# Fast parameterization of Martini3 models for fragments and small molecules

**DOI:** 10.1101/2025.07.13.664596

**Authors:** Magdalena Szczuka, Gilberto P. Pereira, Luis J. Walter, Marc Gueroult, Pierre Poulain, Tristan Bereau, Paulo C. T. Souza, Matthieu Chavent

## Abstract

Coarse-grained molecular dynamics simulations, such as those performed with the recently parametrized Martini 3 force field, simplify molecular models and enable the study of larger systems over longer timescales. With this new implementation, Martini 3 allows more bead types and sizes, becoming more amenable to study dynamical phenomena involving small molecules such as protein-ligand interactions and membrane permeation. However, while there were some solutions to automatically model small molecules using the previous iteration of Martini force field, there is no simple way to generate such molecules for Martini 3 yet. Here, we introduce Auto-MartiniM3, an advanced and updated version of the Auto-Martini program, designed to automate the coarse-graining of small molecules to be used with the Martini 3 force field. We validated our approach by modeling 81 small molecules from the Martini Database and comparing their structural and thermodynamic properties with ones obtained from models designed by Martini experts. Additionally, we assessed the behavior of Auto-MartiniM3-generated models by calculating solute translocation and free energy across lipid bilayers. We also evaluated more complex molecules such as caffeine by testing its binding to the adenosine A2A receptor. Finally, our results from deploying Auto-MartiniM3 on a large dataset of molecular fragments demonstrate that this program can become a tool of choice for fast high-throughput creation of coarse-grained models of small molecules, offering a good balance between automation and accuracy. Auto-MartiniM3 source code is freely available at https://github.com/Martini-Force-Field-Initiative/Automartini_M3

## Introduction

Molecular dynamics (MD) simulations provide insights into motions and flexibility of a diversity of molecules from few atoms to large macromolecular complexes (1–3). Among other applications, they give critical information in drug design (4), by enabling the analysis of thermodynamics and kinetics of ligand binding and unbinding (5, 6), free energy calculations (7), or characterizing protein-ligand interaction pathways (8). The exponential growth of computing ability with graphics processing units (GPUs), exascale supercomputers, as well as dedicated MD software has created exciting new opportunities for modelling large macromolecular assemblies (9–11) such as a whole-virion (12–14) or an entire gene locus (15). Although All-Atom (AA) simulations on such a scale are achievable, they are costly and not easily accessible (16). Thus, lower resolution models, like those based on coarse-grained (CG) methods, can be used for modeling bigger and more complex systems for longer time scales (17, 18).

Among the different CG models available (19–22), the Martini force field is one of the most popular choices. The Martini force field gained its popularity due to its extensive library of models, enabling the coarse graining of lipids and biological membranes (23–26), proteins (27–29), nucleic acids (30, 31), carbohydrates (32), nanoparticles (33) and more. By mixing those components together, it provides hints into behavior of complex molecular systems in a closer-to-reality set up (34). At its beginnings in 2004, this force field was constituted of 4 main building blocks and 8 subtypes (25). Then, in 2007, it was extended to model 18 bead types combined to two bead sizes: regular (R) and small (S) (24). In those earliest versions of the Martini force field, it was difficult to model drug-like molecules because of the limited number of bead types and the 4 non-hydrogen atoms to 1 bead mapping scheme (35). Later on, it was revised by Martini developers, following the parametrization of DNA model (31) and designing a special Tiny bead size dedicated to nucleobases. However, their behavior was the same as the larger regular bead type regarding cross-interactions, introducing some artifacts (36). Since then, numerous updates have considerably increased the diversity of molecular entities, now reaching a vast chemical space with 843 beads, enclosing three sizes of beads and 28 main bead chemical types (34, 37) calibrated against thermodynamic data such as oil/water partitioning coefficients. The release of the Martini 3 force field significantly improved the level of detail and chemical accuracy (37) to model systems such as small molecules and fragments (38), proteins (39), lipid membranes (40), protein lipidations (41), culminating with the modeling of an entire cell (42). With this recent reparametrization, modelling molecular systems containing small molecules is now at reach. Indeed, Martini 3 force field has been employed in recent studies that require precise parametrization of systems containing small, drug-like molecules, like explorations of protein-ligand interactions (43–48), modeling of chemical reactivity (49), drug design (50, 51) and high-throughput pipelines for drug development (51, 52). Nonetheless, small molecule parametrization can be a laborious work, requiring expertise and deep understanding of the chemical properties of the molecules, detailed structure, specific interactions they take part in, and the principles of coarse-grained modeling. Automation of this task is a gateway to efficient use of coarse-grained molecular dynamics simulations in drug discovery pipelines (50, 51, 53). There is a whole Martinidome (34) of programs released to help users create CG models for different kinds of molecules. Examples include Martinize2 (54) for building CG models of proteins, Insane (26) for assembling CG systems, CGCompiler (55), Swarm-GC (56, 57), Bartender (58), or PyCGTOOL (59) for molecule parametrization. However, mentioned tools for ligands coarse graining focus almost exclusively on bonded parameters optimization and need AA simulation trajectories as input data. Other programs that enable complete small molecule parametrization, including mapping and bonded/nonbonded interactions definition, are available (60, 61), but those approaches were developed to work with the Martini 2 force field. Thus, there is a need to develop an automated workflow for designing CG models of small molecules in synergy with the Martini 3 force field.

Here, we introduce Auto-MartiniM3, an updated version of Auto-Martini (60) designed for the automated coarse-graining of small molecules for the Martini 3 force field. It corresponds to a redesign of the original Auto-Martini, here referred as Auto-MartiniM2, which followed the mapping rules of the Martini 2 force field (24). First, we explain the changes made to Auto-MartiniM2, which include updating the mapping algorithm and parametrization to suit the new Martini 3 rules. To test the program, we apply Auto-MartiniM3 on a dataset of 85 small molecules for which human-crafted models exist in the Martini Database (MAD library) (62), manually designed by Alessandri and collaborators (38). We compare our models to this reference set and we simulate them to estimate their numerical stability. To confirm the validity and ensure proper behavior of these new parametrizations, we test the bead assignment by computing solvent partitioning free energies between octanol/water, hexadecane/water, and chloroform/water. Additionally, we also carry out potential-of-mean-force (PMF) calculations for seven small-molecule/membrane systems. Furthermore, we confirm that models created by Auto-MartiniM3 can lead to meaningful results by evaluating the binding of caffeine to the adenosine A_2A_ receptor. Finally, we show that our program can be used in drug development pipelines as a method of choice for rapid, “on the fly” coarse graining of small molecules by setting-up CG models of 100,000 small molecules from the Enamine compounds library (https://enamine.net/compound-libraries).

## Methods

### Strategy

Coarse graining of small molecules needs to take into account their variability, detailed chemical structure and specific interactions they take part in. To tackle those issues, and create a transferable model, the Martini force field adopts a building block approach utilizing a finite set of pre-parametrized bead types. They introduced polar (P), intermediately polar (N), non-polar (C) and charged (Q) beads, as well as additional types describing water molecules (W), divalent ions (D) and groups containing halogens (X). Each bead is characterized by a water/oil partitioning coefficient, meaning they can be classified according to their hydrophobicity, which influences how they interact via Lennard-Jones (LJ) potential. In the Martini 3 version the range of available bead sizes is expanded with the use of tiny (T) beads, apart from small (S), and regular (R) beads, grouping respectively, in average, 2, 3, or 4 non-hydrogen atoms and corresponding hydrogens into a single bead, with LJ parameter σ of 0.34, 0.41 and 0.47 nm (37). These changes enable a detailed mapping of rings— fragments which are recurrent in small molecules. For instance, to create accurate Martini 3 models, it is advisable to use T beads to map aromatic rings and S beads for aliphatic rings, which helps reflect the bulkiness of those structures (38). Functional groups should be kept intact whenever possible, to preserve their chemical identity and interactions (37). For establishing bonded parameters, it is recommended that the center of beads should be treated as a center of geometry of the atoms (including hydrogens) and not as a center of mass, which was commonly used in the previous version of the Martini force field for certain classes of molecules such as proteins (27).

Bead assignment strategies described above are inspired by a top-down approach, implemented in Martini, which involves deriving non-bonded interaction parameters by fitting them to experimental data, ensuring that the model accurately reflects real, macroscopic properties. For bonded parameters, a bottom-up approach (63) is usually applied to extract parameters from detailed atomistic simulations and implement them in the resulting coarse-grained model, allowing the force field to capture molecular-level interactions with high fidelity. Combining these approaches and rules ensures that Martini 3 balances accuracy and computational efficiency across a wide range of systems. Thus, taking into account all the recent changes and implementations of Martini 3 and turning them into a set of algorithmic rules for automatic parametrization is not straightforward and requires compromises, which determines the core of the Auto-MartiniM3 algorithm. The benefit of using Auto-Martini is its rapidity and the use of one simple input required to run the program, making it a perfect candidate for managing large amounts of data, for example in high-throughput screenings (64). As in its original version from 2015 (60), the algorithm distributes the closest atoms into beads. Then, it defines bonded parameters and bead types, and produces a topology and coordinate files, ready to be used in an MD simulation. The Auto-MartiniM3 algorithm follows overall the same procedure as presented in **Figure 1**. In the next sections, we will describe in detail the changes in the workflow to determine the AA to CG mapping, bonded parameters, and the assignment of bead types and sizes for nonbonded interactions in agreement with the Martini 3 force field rules for small molecule mapping (38).

**Figure 1.**
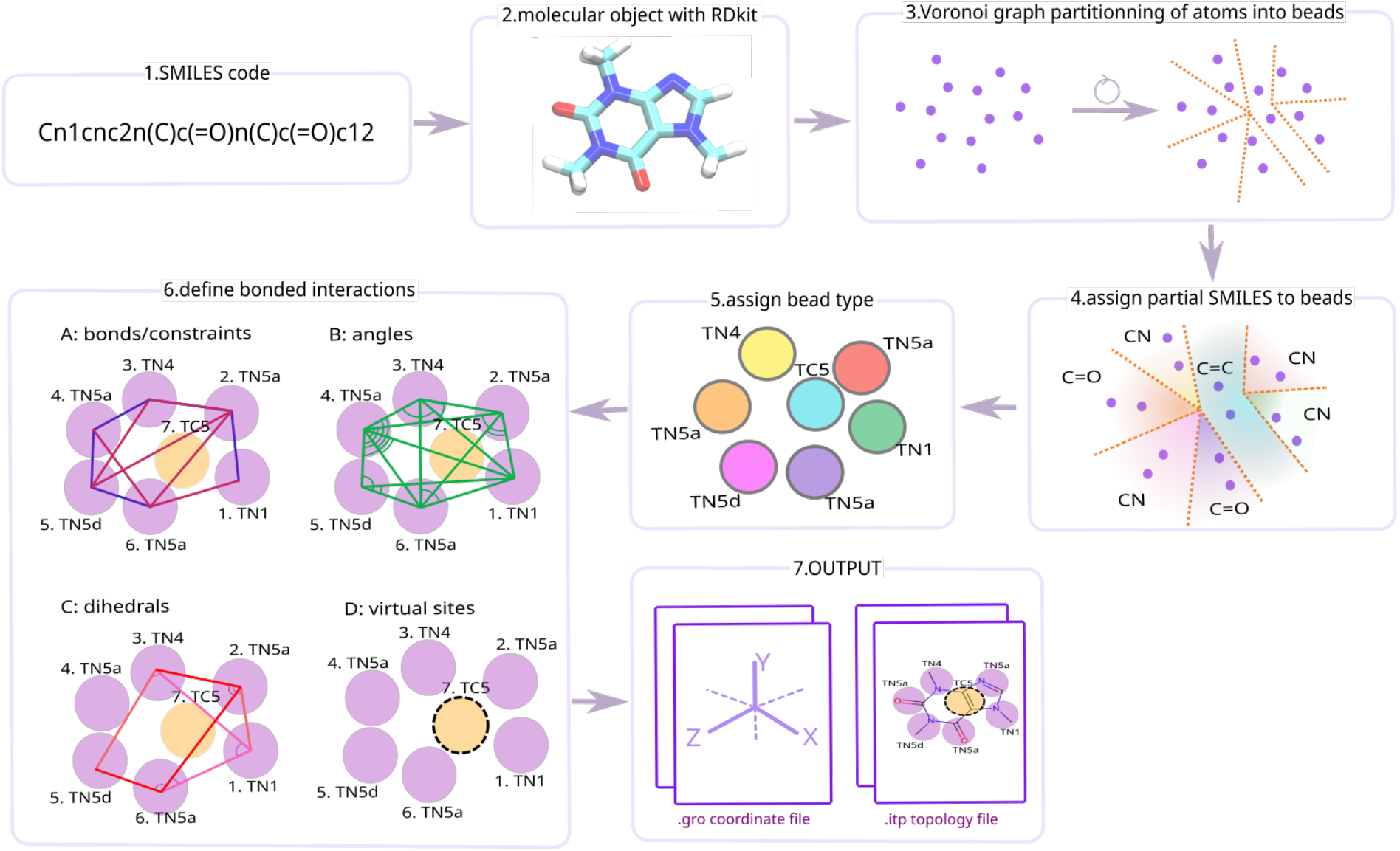
Overall pipeline of Auto-MartiniM3 algorithm. Parametrization workflow example for the caffeine molecule. The algorithm takes as input the SMILES code of a molecule **(1)** and transforms it with RDkit into a 3D molecule object with coordinates assigned to each atom **(2)**. Coordinates are used to assemble two to four closest non-hydrogen atoms into beads **(3)** with assigned partial SMILES code, representing enclosed atoms **(4)**. Partial SMILES code is then used to define bead type, taking into account the physicochemical behavior of each bead in water/oil solvent **(5)**. Furthermore, every bead’s center of geometry is used to define bonded interactions, such as bonds **(6A)**, angles **(6B)**, dihedrals **(6C)** and virtual sites **(6D)**. Bonded and nonbonded parameters are saved in the topology file and the coordinates of the bead are saved in the coordinates file **(7)**, ready for molecular dynamics simulation with GROMACS.

### Mapping

The parametrization starts with a simple input: the SMILES code of the molecule, which describes the structure as a string of characters (65). Coarse graining of a molecule requires its spatial image in order to recreate the intramolecular interactions, therefore, like in the previous version, Auto-MartiniM3 uses the RDKit Python package (66), to transform the SMILES code into a 3-dimensional molecular object, which is further analyzed (**Figure 1-(1,2)**). The purpose of mapping is to describe a structure of a given molecule by assigning an optimal set of coarse-grained beads in place of the atoms. This assignment is done with the Voronoi graph partitioning algorithm, present in Auto-MartiniM2, which groups together between 2 and 4 atoms per bead (**Figure 1-(3)**). The algorithm generates all possible mapping options for the molecule within this scope, leading to an exponential increase in the number of partitions as the number of non-hydrogen atoms grows. For each possible partition, the energy function is assigned, which quantifies the pertinence of a given mapping, thus enabling the ranking of the mappings and identifying the one with most optimal energy (60). In Auto-MartiniM3, values of energy function parameters are adapted to force the use of newly added Tiny beads, more suitable for parameterizing small molecules.

In order to keep the integrity of functional groups within molecular structures, we have adapted the RDKit Ertl Functional Groups Finder algorithm to recognize functional groups within a given molecule and accept the mappings which encapsulate them in one bead (52). It influences the assignment of bead types which improves the physicochemical description and behavior of a given molecule. In cases where it is impossible to keep a whole functional group within a bead, for example due to the number of atoms involved, the algorithm reruns the mapping and allows for partial conservation of the functional group. Another new feature introduced in Auto-MartiniM3 is the use of the center of geometry of every bead as a bead coordinate. In Auto-MartiniM2, every CG bead was placed on one of the N atoms. Then, the atoms closest to a given bead were grouped together to form a CG bead, in accordance with the score of the energy function. The center of bead was alas neither the center of mass nor the center of geometry, but it was defined with the coordinates of an atom, upon which the CG bead was firstly placed. In the updated version, the center of each bead is recalculated by taking into account the coordinates of every atom, including hydrogens. Finally, for each set of atoms, the algorithm determines a partial SMILES code which serves as an input to find an octanol/water partitioning coefficient (logP) (see details in the nonbonded interactions subsection). This value allows one to find the most suitable bead type, following the guidelines of Martini 3 force field developers (38).

### Nonbonded interactions

The parametrization of nonbonded interactions is challenging, because it ought to reflect the collective behavior of a group of atoms. Martini 3 force field, with its building block approach, handles this by giving a vast selection of 843 bead types (34) and establishing interaction parameters between each possible pair, allowing more extensive mapping possibilities. As described before, this force field update includes polar, intermediately polar, non-polar and charged types of beads, with additional labels, as hydrogen donor/acceptor, to reinforce complementary hydrogen bonds, as well as beads containing divalent ions and halogen atoms. Each type of bead can be described with the value of the partitioning free energy between polar and apolar phases, for example water/octanol coefficient (logP), which is available in the Supporting Information of Martini 3 force field publication (37). In Auto-MartiniM3, this coefficient is assigned for each bead by associating partial SMILES, representing grouped atoms of a given bead, to preliminary calculated octanol/water partitioning coefficients defined in our custom database file (https://github.com/Martini-Force-Field-Initiative/Auto-martini_M3/blob/main/auto_martiniM3/logP_smi.dat). If a particular representation is not found in this file, the ALOGPS program (67) is used to find corresponding logP coefficients, as done in Auto-MartiniM2. With associated logP value to a given partial SMILES, the algorithm searches for the nearest water/octanol partitioning coefficient of Martini3 beads, and selects an appropriate bead type and size (**Figure 1-(5)**).

### Bonded interactions

The assignment of bonded interactions uses the center of geometry of every bead. To obtain accurate models in terms of volume and distances between the beads, we mimic an MD minimization using the RDKit package on an AA molecular object created with this tool. This minimization is implemented before coarse-graining to slightly “relax” the molecular structure, making the 3-dimensional molecular model more accurate in reproducing correct distances between atoms, and ultimately beads. Besides this change, the determination of the CG topology is similar to the previous version of the program, which considers that any CG bead *i* is connected to *j* via bond, if an atom from *i* is chemically bonded to an atom from *j*. The angles and improper dihedrals are constructed from the collection of bonds and their values are calculated with new definitions of bead centers. Bonded interactions are restrained using harmonic potentials, based on data from state-of-the-art mappings made manually by Martini3 developers, available in MAD (Martini Database) (62). According to the guidelines for small-molecule parametrization in Martini 3 force field (38), fused rings should be held by constraints in order to preserve the rigidity of the structure. The use of stiff potentials requires a short time step, which can be computationally expensive. To mitigate this issue, the use of virtual sites can preserve numerical stability without enforcing shorter time steps. To do so, the algorithm places a virtual site whenever a structure contains fused rings (**Figure 1-(6)**). It finds the most connected bead, which is the bead with the largest number of bonds, and transforms it into an N-body virtual site. For molecules with only one ring and chemical groups attached to it, besides other bonded parameters, we applied exclusion on the most distant pair of beads, where one bead is a ring bead and the other is a non-ring bead, attached to this ring (see for example benzotrifluoride (BTZF), in **Figure S1**).

### Simulation parameters

#### CG simulations of molecules to compare with MAD molecules

To compare with the state-of-the-art modeling of small molecules, we model 81 molecules (**Figure S1**) using Auto-MartiniM3 and subsequently perform short simulations of these molecules in a water box with GROMACS 2023.2. For minimization, we use the steepest descent algorithm with the LINCS constraints algorithm. Equilibration is done in three stages, starting with an integration step of 5 fs, followed with 10 fs, and finishing at 20 fs, for 10 ns of simulation at each stage. The Verlet algorithm is used as a cut-off scheme with the straight cut-off at 1.1 nm. The temperature is controlled using the velocity-rescale (68) algorithm and set at 310 K. The pressure is controlled using the Berendsen barostat (69) with a compressibility of 3 · 10^−4^ bar^-1^, a coupling parameter of 4.0 ps, and an isotropic pressure of 1 bar. Then, a production run is performed for 100 ns, applying a 20 fs integration step. The velocity-rescale thermostat coupled with the Parrinello-Rahman barostat (70) provide control over the temperature, set at 310 K, and an isotropic pressure, set at 1 bar with a compressibility of 3 · 1^−4^ bar^-1^ and a coupling parameter of 4.0 ps.

#### Solvation free energy simulations

To evaluate the quality of the models produced by Auto-MartiniM3, we compute the transfer free energies (ΔΔG) of each model between water and three organic solvents used as gold standard in Martini 3 parametrization: hydrated octanol (OCT), chloroform (CLF) and hexadecane (HD) (**Figure 3**). These calculations are used to validate the performance of Auto-MartiniM3 against experimental data, as done for previous work (38). Thus, for each model 4 sets of thermodynamic integration simulations are set up, each with the goal of computing the solvation free energy of a given model in one of the four solvents (ΔG_vac-W_, ΔG_vac-OCT_, ΔG_vac-HD_, ΔG_vac-CLF_, with vac meaning vacuum). Each system is first solvated into the corresponding solvent in a 5.5×5.5×5.5 nm^3^ simulation box. In the case of hydrated octanol, 20% water is included in the simulation box as observed experimentally for biphasic water/octanol partitioning systems (71). Systems are energy minimized using the steepest descent algorithm for a maximum of 10,000 steps with LINCS being used to control constraints. A short equilibration step follows, for 5 ns with an integration timestep of 10 fs and with the Verlet algorithm as a cut-off scheme (straight cut-off at 1.1 nm). The temperature is controlled using the velocity-rescale (68) algorithm and set at 298K and pressure is controlled using the Berendsen barostat (69) with a compressibility of 3 · 10^−4^ bar^-1^ and a coupling parameter of 4.0 ps at 1 bar pressure. The solvation free energies are computed using thermodynamic integration, an alchemical free energy method. For each system, a series of 11 simulations with equally spaced λ windows are carried out with a stochastic dynamics integrator with a 20 fs timestep and employing the Parrinello-Rahman barostat (70) and a compressibility at 1 bar with coupling parameter of 4.0 ps. Each window runs for 5 ns, using a soft-core potential while setting *α* = 0.5 and power set to 1. The Bennet Acceptable Ratio (72) is used to compute the free energies and the associated error. The free energy of transferring a given molecule from water to one of the other solvents (OCT, HD, CLF) is computed as the difference between the solvation free energy of that molecule in each solvent (ΔΔG_OCT-W_ = ΔG_vac-OCT_ - ΔG_vac-W_) (73). Simulations are carried out using GROMACS 2023.2 (74, 75).

#### Potential of Mean Force of molecule insertion in lipid membrane

To further assess the quality of the parametrization, we compute the potential of mean force (PMF) of Auto-MartiniM3-generated molecules inserted into phospholipid bilayers. The PMFs are compared with results from Auto-MartiniM2 (60) and atomistic references. The preparation and simulation of the bilayer systems are based on the protocol proposed by (40). The phospholipid bilayers are generated and solvated by the program insane (26) using a leaflet area of 6 nm x 6 nm. Subsequently, the bilayer system is energy minimized and equilibrated for 2 ns with an integration timestep of 20 fs. We use umbrella sampling (76) to calculate the PMF of the molecule insertion. The collective variable employed is the distance to the bilayer center, projected on the bilayer normal. To improve the efficiency of the simulations and leverage the bilayer’s inherent symmetry, two identical molecules are introduced into the membrane, following the approach previously described by (77). This yields 92 umbrella windows, with a distance to the bilayer center between 0 and 3.95 nm. To improve simulation stability within the lipid bilayer, we reduced the angle and dihedral force constants suggested by Auto-MartiniM3 for three ligands. Specifically, dihedral constants were set to 0.5 kJ/mol for dibenzantracene, ibuprofen, and alprazolam, with alprazolam also having its angle force constants reduced by a factor of 100. As these modifications do not substantially affect ligand conformations, we do not expect a significant impact on the resulting PMF profiles. Each umbrella system is energy minimized using a steepest descent algorithm. This step is followed by two equilibration simulations of 100 ps and 2 ns with integration timesteps 10 fs and 20 fs, respectively. Finally, each umbrella window is subjected to a 120-ns production simulation with a 20-fs timestep and a harmonic biasing potential of 1000 kJ/mol/nm^2^. The temperature of the system is maintained at 297K using the velocity-rescale thermostat (68). The pressure is set to 1 bar and controlled using a semi-isotropic Parrinello-Rahman barostat (70) with a compressibility of 3 · 10 ^−4^ bar^-1^. Regarding the non-bonded interactions, we employ a cutoff of 1.1 nm in conjunction with a potential shift. For electrostatic interactions, a reaction field with a dielectric constant of 15 is used. To prevent membrane artifacts (78) we manually set the rlist parameter to 1.35 nm. The simulations are carried out using GROMACS 2024.2 (75). To obtain the final PMF profiles, we use the weighted histogram analysis method (WHAM) (79).

#### Interaction of caffeine with A2A receptor embedded in POPC membrane

The molecular dynamics simulations are performed in GROMACS 2023.2 (74, 75) and simulations set ups are created by following the work of Souza and co-authors (47). For minimization, we use the steepest descent algorithm with the LINCS constraints algorithm. System is equilibrated in four stages, with integration steps of 2 fs, 5 fs, 10 fs and finishing with an integration step of 20 fs, for 10 ns at each stage and with the Verlet algorithm as a cut-off scheme with the straight cut-off at 1.1 nm. The temperature is controlled using the velocity-rescale algorithm (68) and set at 320 K. The pressure is controlled using the Parrinello-Rahman barostat (70) with a compressibility of 3 · 10 ^−4^ bar^-1^ and a coupling parameter of 12.0 ps, and a semi-isotropic pressure at 1 bar. We run 12 production runs of 20 microseconds (**Figure S7**), applying a 20-fs integration step using the same thermostat and barostat algorithms as done in the equilibration step.

#### Data processing, analysis, and availability

Simulated trajectories were analyzed with in-house Python scripts based on NumPy (80), Matplotlib (81), SciPy (82) and MDAnalysis (83). Visual molecular dynamics (VMD) version 1.9.3. (84) was used for image generation and visual inspection of the simulated systems. The calculation of spatial properties is carried out by using GROMACS 2023.3 (74, 75) commands using gmxsasa − ndots 4800 − probe 0.191 to take into account CG bead size for SASA calculations. We have used GROMACS functions gmxrms and gmxrmsf for RMSD and RMSF respectively.

All the data produced during the simulations, as well as Auto-MartiniM3 models are available in Zenodo (https://doi.org/10.5281/zenodo.15783892).

#### Source code availability

Auto-MartiniM3 source code is available on GitHub (https://github.com/Martini-Force-Field-Initiative/Automartini_M3) and archived in Software Heritage (swh:1:dir:24f92272c68221b9d12e7cf7c2e8e97e67547657)

## Results

### Comparison of State-of-the-art and Auto-MartiniM3 Coarse-grained Parametrizations

To first test the program, we use 85 SMILES codes of small molecules available in the Martini Database (MAD) (62) from the R. Alessandri dataset (38) to create models with Auto-MartiniM3 and simulate each molecule in a box of water molecules. We manage to model 81 out of 85 molecules (**Figure S1**). Some molecules could not be fully parameterized with Auto-MartiniM3 due to the presence of certain functional groups, such as nitro groups, which are not supported yet by the ALOGPS algorithm. Overall, we simulate 78 molecules in a stable and reproducible manner, and the quality of parametrization is evaluated by comparing Auto-MartiniM3 models with state-of-the-art models from MAD, as shown in **Figure 2**. Looking at the number of molecules with a given bead’s nature, Auto-MartiniM3 produces more mappings with polar beads (P) and less mappings with intermediate beads (N) than the manual mappings in MAD, suggesting that Auto-MartiniM3 tends to yield more hydrophilic molecules. The similar high number of non-polar beads (C) in both Auto-MartiniM3 and MAD models can be attributed to the presence of aromatic cycles in small molecule structures, which are modeled using apolar TC5 beads. Overall similar distribution of bead types per molecule as well as bead size (**Figure S2**) proves that Auto-MartiniM3 parametrization is in agreement with manual parametrizations made by experts in the field.

**Figure 2.**
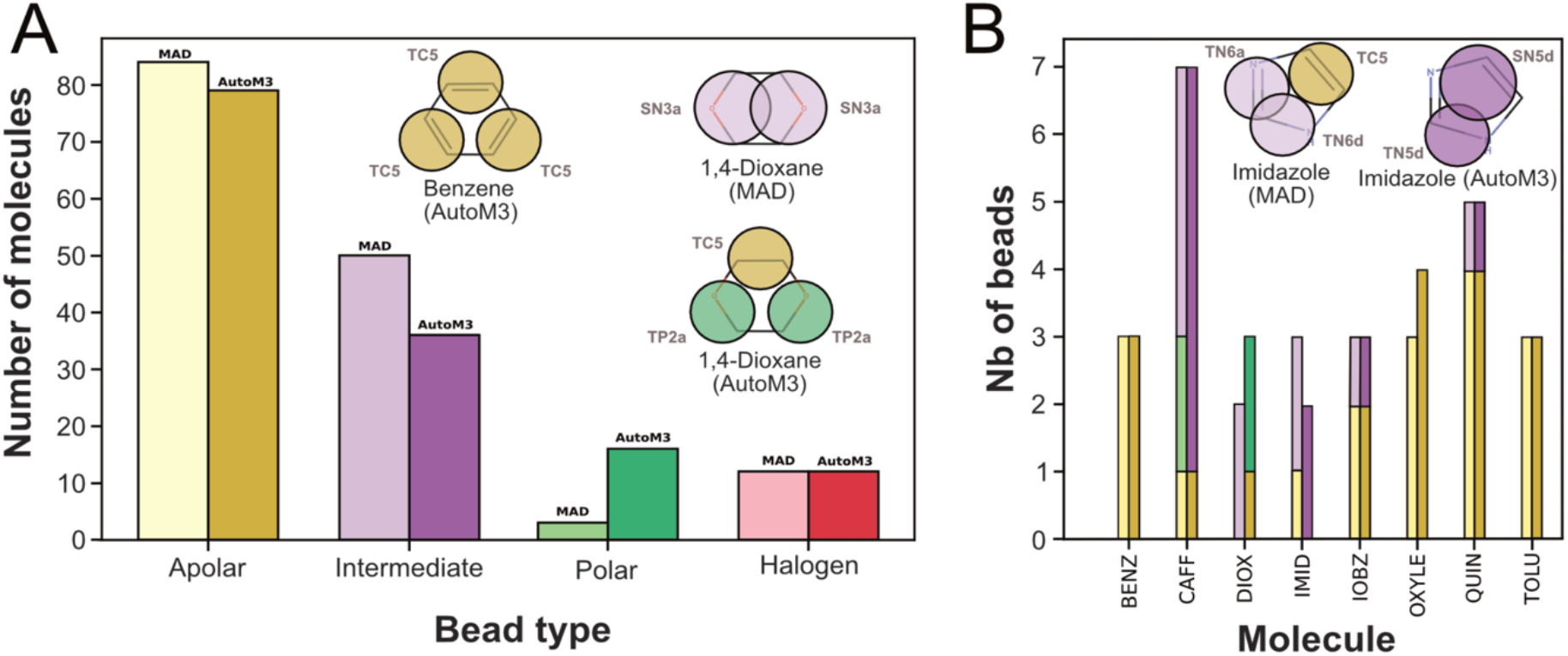
Comparison of topologies created by Auto-MartiniM3 and Martini3FF developers. **(A)** Number of molecules containing at least one bead type extracted in topology files created by Auto-MartiniM3 and by Martini 3 force field developers, available in the Martini Database (MAD) (62). **(B)** Examples of parametrizations made by Martini 3 developers (left) and by Auto-MartiniM3 (right) presented for comparison: benzene (BENZ), caffeine (CAFF), 1,4-dioxane (DIOX), imidazole (IMID), iodobenzene (IOBZ), o-xylene (OXYLE), quinoline (QUIN) and toluene (TOLU) molecule models. Bead types are differentiated by colors, apolar beads in yellow, halogen beads in red, polar beads in green, and intermediate polarity beads in purple. Representations of the 81 models are available in **Figure S1**.

### Spatial Accuracy: Insights from SASA Analysis, RMSD and RMSF Calculations

By comparing solvent-accessible surface area (SASA) metrics from coarse-grained simulations with those obtained from AA structures, we can assess how accurately the CG models capture essential structural features. SASA obtained with Auto-MartiniM3 models reproduce correctly those from AA references, even if there are some Auto-MartiniM3 models with slightly underestimated SASA (**Figure 3-A**). For example, chlorpropham (CLPR), 1,2-dichlorobenzene (DCLBZ), N-boc-2-aminophenol (NBAPH) and p-cymene (PCYM) models include regular beads, which reduces the number of beads, thus decreasing SASA by 0.6-0.9 nm^2^. In the case of anthracene (ANTH) and 2,2’-bithiophene (2T), models generated by Auto-MartiniM3 have less beads than state-of-the-art models. 2T Auto-MartiniM3 model has 4 beads (TC5-SC6-SC5-TC6), while the MAD model is constituted by 6 beads (with two being virtual sites), which, again, may explain the decrease of SASA. This may arise from a limitation of Auto-MartiniM3: the mapping does not allow bead sharing of atoms, which may help to get the right symmetry for certain molecules. In this case Auto-MartiniM3 may parameterize one small bead instead of two tiny beads. Nevertheless, when comparing with the original Auto-MartiniM2 parametrization (**Figure 3-C**), it is clear that the agreement between the references and Auto-Martini3 models is enhanced due to the addition of tiny beads in Martini 3. Furthermore, the bead-bead distances generated by Auto-MartiniM3 are more broadly distributed than in Auto-martiniM2 models (**Figure S3**), which allows a better representation of molecular volume.

**Figure 3.**
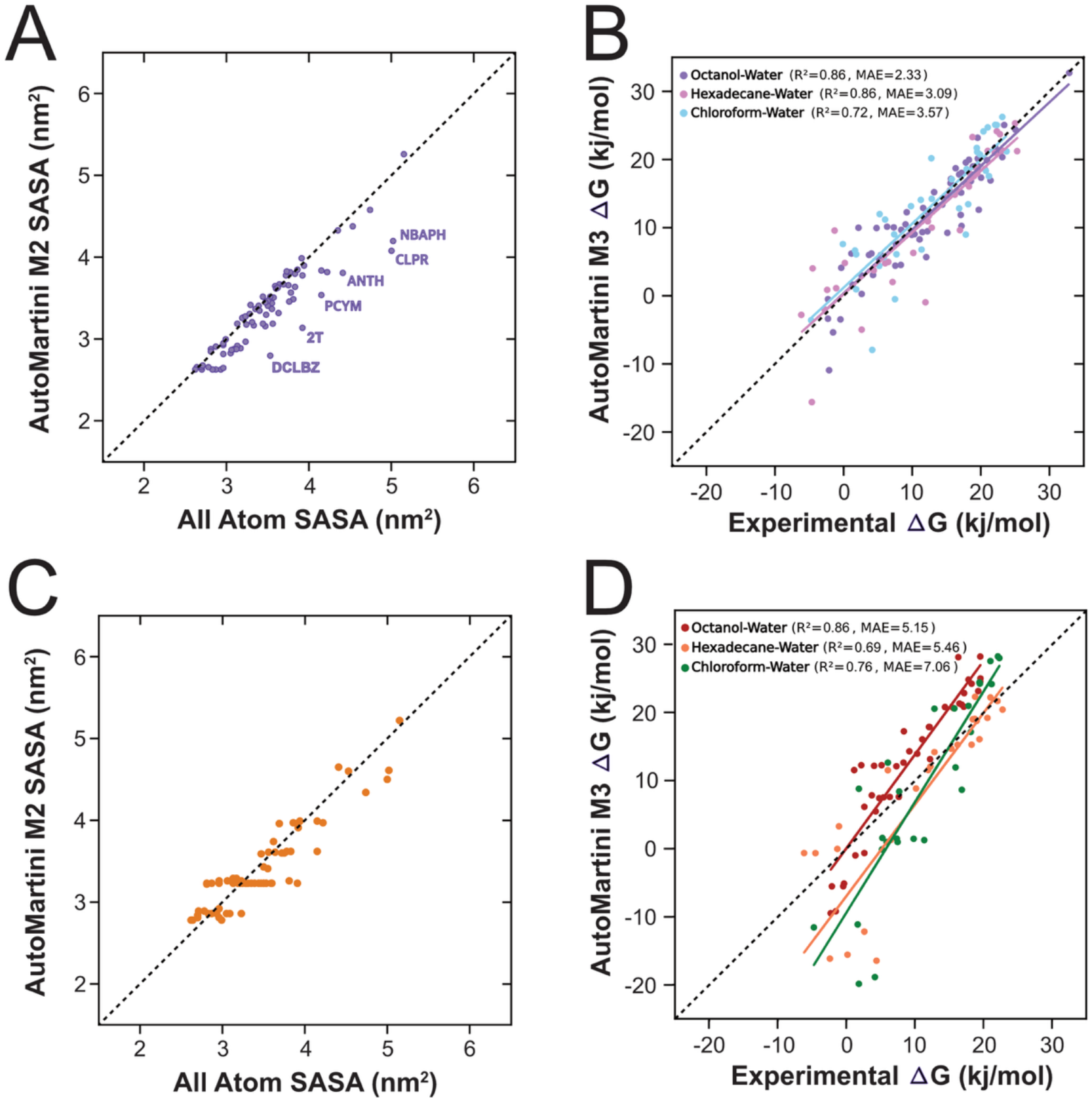
Solvent-Accessible Surface Area and Partitioning free energies in water/oil phases in regard to reference data. SASA of molecules parameterized with our Auto-MartiniM3 **(A)** and the original Auto-MartiniM2 **(C)** in comparison with AA models. Thermodynamic Integration calculations based on models generated by Auto-MartiniM3 **(B)** and Auto-MartiniM2 **(D)** plotted against experimental data. Labels in (A) correspond to molecule presented in Figure S1.

We also perform root-mean-square deviation (RMSD) and root-mean-square fluctuation (RMSF) analyses to assess the overall stability of the coarse-grained models generated by Auto-MartiniM3. Our results suggest that, overall, Auto-MartiniM3 is capable of generating coarse-grained models for small molecules that maintain structural stability (**Figure S4**). In some cases, molecules may exhibit somewhat high RMSD and RMSF values (**Figure S4**). To further stabilize the molecules, increasing forces for bonded parameter, angles, and improper dihedrals may solve this issue (see for example DMBZQ molecule **Figure S4**).

### Water-oil Partitioning Behavior

We then benchmark the bead assignment of the Auto-MartiniM3 parametrization scheme by comparing calculated water/oil logP with experimental data used as reference in the Alessandri dataset (38). As in the original publication, we compute free-energy changes associated with the transfer of small molecules between water and octanol, hexadecane and chloroform phases, respectively. The partitioning free energy values from Auto-MartiniM3 are in agreement with experimental data, exhibiting mean absolute errors of 2.33, 3.09, and 3.57 kJ mol^-1^ for octanol-water, hexadecane-water and chloroform-water, respectively (**Figure 3-B**). There is a clear improvement over the original version (**Figure 3-D**), which yields mean absolute errors of 5.15, 5.46, and 7.06 kJ mol^-1^ for octanol-water, hexadecane-water and chloroform-water, respectively. The same data plotted against results from the literature (Alessandri et al., 2022) (**Figure S5**) shows that Auto-MartiniM3 produces models close in quality to those generated by Martini 3 developers.

### Free-energy Profile of Insertion into a Lipid Membrane

We further evaluate the Auto-MartiniM3-generated models in a solvent-membrane environment by calculating the potential of mean force (PMF) along the distance to the bilayer midplane z for seven different molecules. We evaluate 4-ethylphenol, 5-phenylvaleric acid, ibuprofen, alprazolam and theophylline in a 1,2-dioleoyl-sn-glycero-3-phosphocholine (DOPC), propanol in a 1,2-dilauroyl-sn-glycero-3-phosphocholine (DLPC) membrane, and dibenz[a,h]anthracene in 1-palmitoyl-2-oleoyl-sn-glycero-3-phosphocholin (POPC) membrane. The resulting PMFs are compared to AA references from (85, 86), and the original Auto-MartiniM2 publication (60) (**Figure 4**). This analysis aims to validate the Auto-MartiniM3 models in terms of structural thermodynamics of a small molecules at complex membrane interfaces.

**Figure 4.**
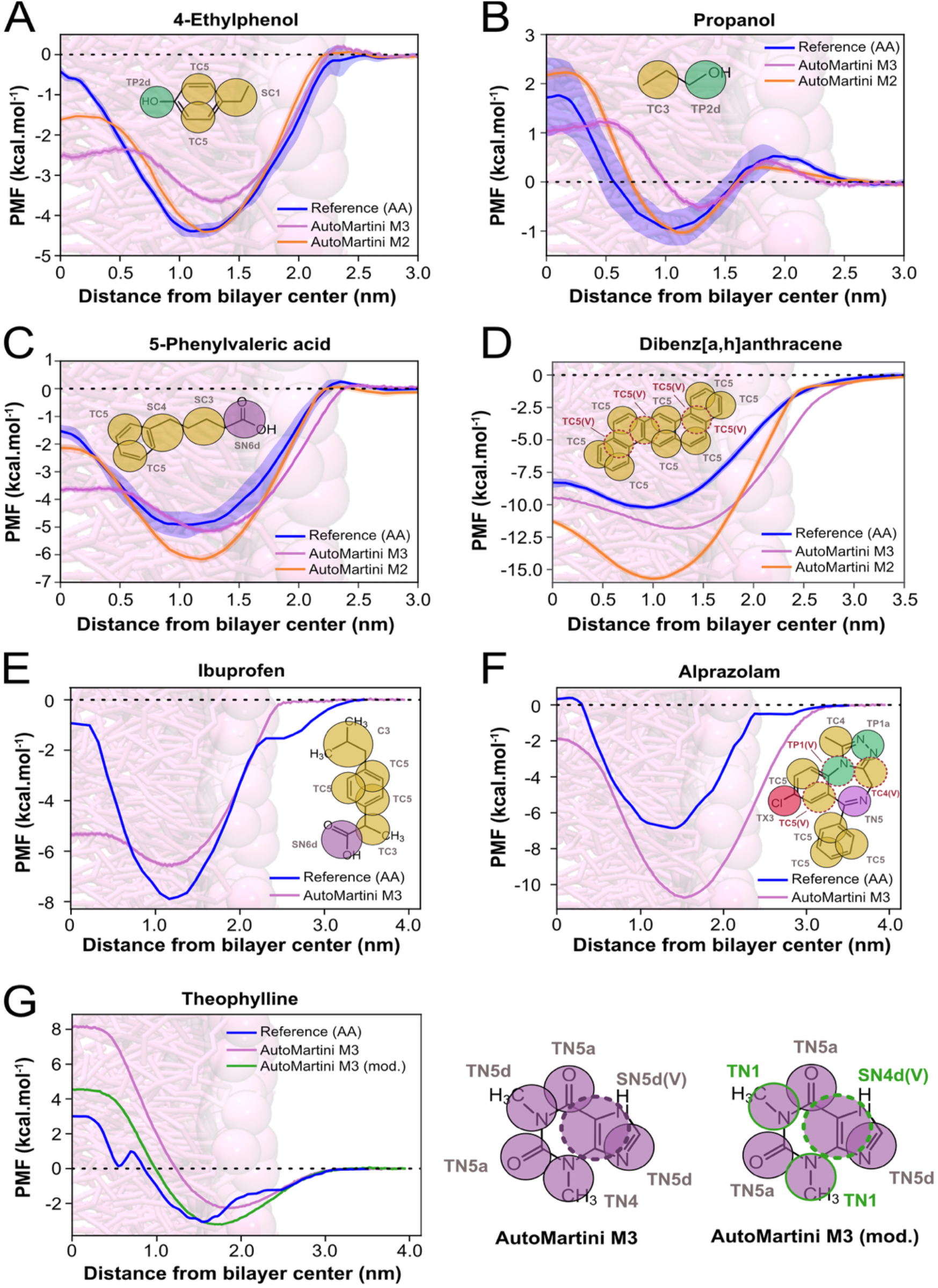
Potential of Mean Force of molecule insertion in lipid membrane. **(A)** 4-ethylphenol in DOPC/water, **(B)** propanol in DLPC/water, **(C)** 5-phenylvaleric acid in DOPC/water, **(D)** dibenz[a,h]anthracene in POPC/water, **(E)** ibuprofen in DOPC/water, **(F)** alprazolam in DOPC/water, **(G)** theophylline in DOPC/water. In green manual modifications of some bead types for theophylline parametrization. PMF of molecules parameterized with Auto-MartiniM3 are in pink, data of molecules from original Auto-MartiniM2 paper in orange and AA reference data in blue. Atomistic and Auto-MartiniM2 data are extracted from (60) (A-D) while AA references presented in (E-G) are from from (85).

As expected, the coarse-grained PMFs are generally smoother and exhibit more gradual transitions compared to their atomistic counterparts, reflecting the reduced resolution of the force field. Nevertheless, the insertion profiles of all Auto-MartiniM3 models closely follow the AA references, with energy minima located at comparable positions. The agreement between Auto-MartiniM3 and atomistic PMFs is similar with that of Auto-MartiniM2. For instance, dibenz[a,h]anthracene shows an improved match in Auto-MartiniM3 (**Figure 4D**: RMSD of 2.1 kcal/mol) compared to Auto-MartiniM2 (3.6 kcal/mol), whereas 4-ethylphenol **(Figure 4A**: RMSD of 0.8 kcal/mol vs. 0.4 kcal/mol), propanol (**Figure 4B**: 0.6 vs. 0.4 kcal/mol), and 5-phenylvaleric acid (**Figure 4C**: 0.8 vs. 0.7 kcal/mol) show slightly higher deviations. It is important to note that AA PMF references available in the literature are limited, and their validation is non-trivial—introducing some uncertainty when using them as benchmarks for coarse-grained models.

For ibuprofen (**Figure 4E**), coarse-grained and atomistic PMFs align well for z > 0.7 nm but deviate near the bilayer center by ∼4 kcal/mol, suggesting a slightly too lipophilic bead type assignment (overall RMSD of 1.7 kcal/mol). More substantial discrepancies are observed for alprazolam and theophylline (**Figure 4F–G**), with RMSDs of 2.9 and 3.2 kcal/mol, respectively. These deviations highlight the challenge of accurately predicting the collective behavior of multiple coarse-grained beads from individual atomistic fragments in larger molecules (e.g., alprazolam, represented by 11 beads). Such cases may require manual refinement, as illustrated for theophylline, where adjusting a few bead types significantly reduce the RMSD to 1.2 kcal/mol (**Figure 4G**). We anticipate that expanding and refining the underlying logP-based fragment database (see Nonbonded interactions section) will minimize the need for such adjustments in future iterations.

### Protein-Ligand Interaction Test Case: Binding of Caffeine Molecule to A_2A_ Receptor

A proof-of-concept illustrating Auto-MartiniM3 applicability in protein-drug interaction studies can be established by simulating the A_2A_ receptor (PDB code: 3RFM) embedded in a POPC membrane. The A_2A_ receptor plays a crucial role in mediating various physiological processes, such as neurological signaling, which makes it a well-known target for drugs treating conditions like Parkinson’s disease (87). Thus, this receptor is a valuable model for studying ligand-receptor interactions and testing new therapeutic compounds. One of the well-known ligands for this receptor is caffeine, which acts as an antagonist and has a highly stimulating effect (87, 88). Moreover, caffeine has its various applications in different fields of medicine (88). For our test case, the caffeine model is coarse grained using Auto-MartiniM3 and ten molecules of caffeine are positioned in the solvent environment (**Figure 5-A**), as done previously (47). Over the course of the 12×20µs simulations (**Figure S7.1-12**), the caffeine molecules are monitored to evaluate their ability to locate and bind to the receptor’s binding site. The results show that the ligand successfully binds to the A_2A_ binding site (**Figure 5-C**). The RMSD of one caffeine, compared to the coarse-grained crystallographic structure position in the binding site (47) as reference, is depicted in **Figure 5-B**. The Auto-MartiniM3 model of caffeine is able to sample positions close to the binding site identified by crystallography while also exploring from solvent to membrane and receptor (**Figure S6**). We found numerous cases in the 12 simulations performed, where the caffeine explored the crystallographically determined A2A binding site, with an RMSD below 6 Å (**Figure S7.1-12**). Overall, we observe at least 65 binding-unbinding events inside or very near the binding pocket, with different binding pathways consistent with findings from (89), lasting from a few nanoseconds to 1-3 microseconds. Some of these pathways pass through the lipid membrane, which is consistent with previous findings (90).

**Figure 5.**
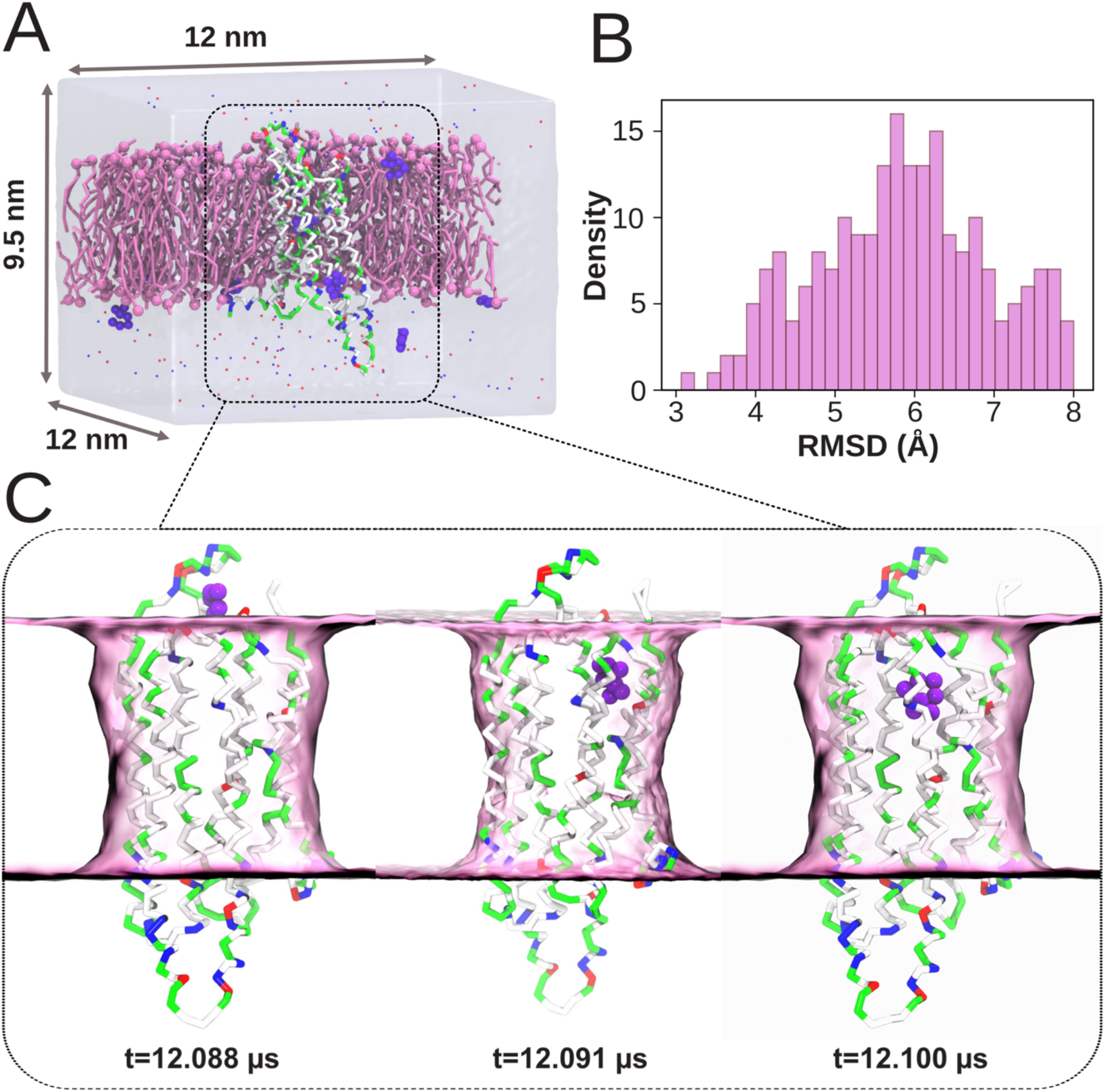
Simulations of Auto-MartiniM3-generated caffeine binding to A2A receptor in POPC membrane. **(A)** System set up with A_2A_ receptor embedded in a POPC (pink) membrane and surrounded by ten caffeine molecules (purple) following the work of (47). **(B)** Histogram of the RMSD (for values under 8Å) of one caffeine during the simulations compared to crystallographic position. **(C)** Example of the entry of a caffeine molecule into its binding site. Solvent ions, lipids and other 9 caffeine molecules are not shown for clarity.

### High-throughput parametrization

We deploy Auto-MartiniM3 on a large dataset (100,000) of molecular fragments (Enamine Fragment Collection) on a supercomputer as previously done with Auto-MartiniM2 (64). Models were generated for 97899 out of 100k molecules, yielding a 97.9 % success rate. Overall, the parametrization experiment took 135 min on high-performance computing resources (2,560 cores, running up to 40 jobs per core at the same time). Detailing the computing time for each molecule (**Figure 6**) reveals a large range of times from less than a second for compounds composed of few heavy atoms up to more than 700 seconds for large compounds constituted of 20 heavy atoms. In average parametrization of a molecule takes around 30 seconds.

**Figure 6.**
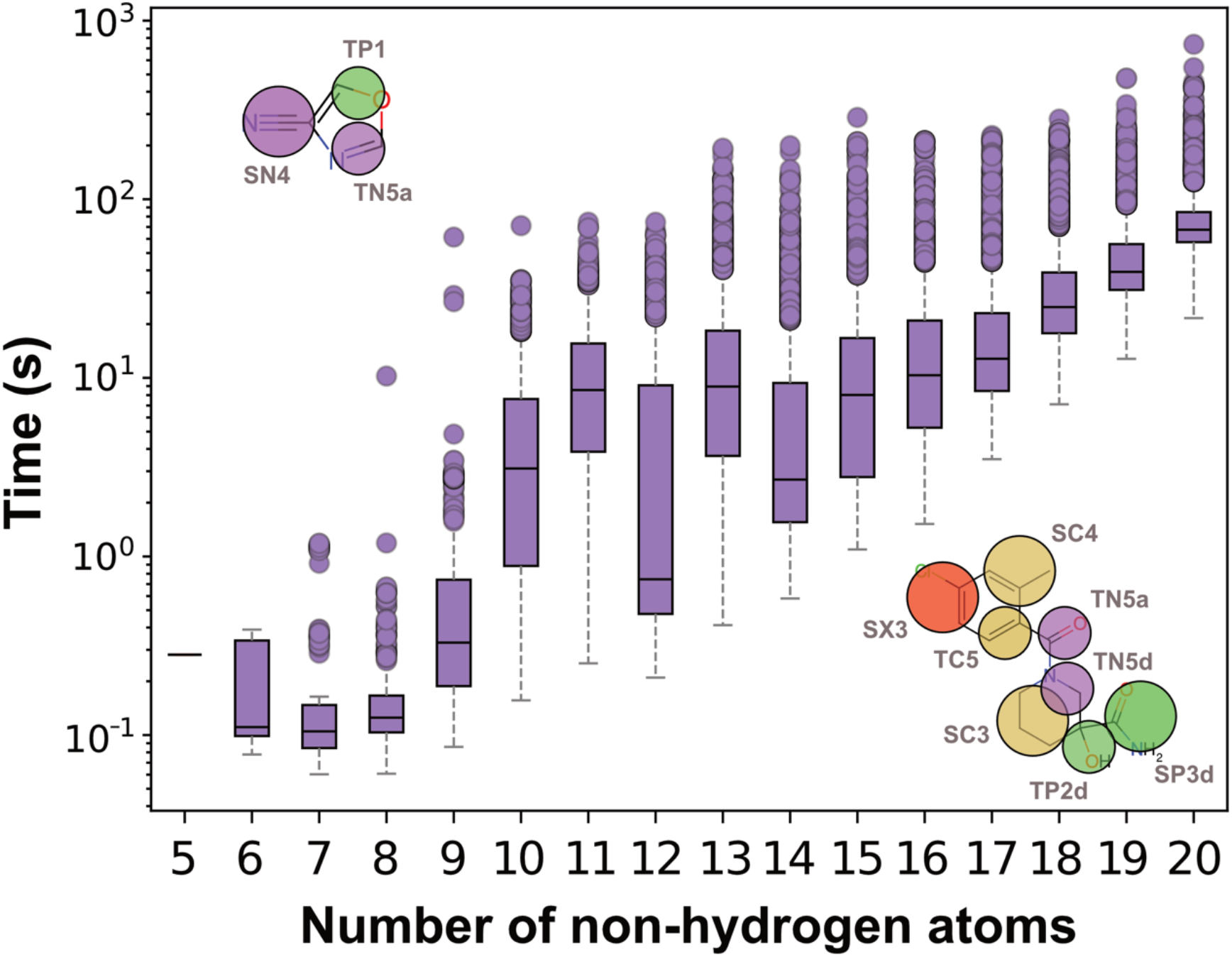
Performance of Auto-MartiniM3. Parametrization time of 100 000 molecules of different sizes from Enamine Fragment Library, parameterized with Auto-MartiniM3. Top left, Auto-MartiniM3 parametrized the Z1143441220 compound in 0.6 s. Bottom right, Auto-MartiniM3 parametrization of the Z2061576810 compound done in 737 s.

## Discussion

Parameterization of molecules, fragments, and other system components is, when needed, typically one of the most challenging steps in setting up a Martini simulation. These parameterization efforts are, more often than not, time-consuming and prone to error, as users must develop mapping schemes and bead assignments based on experience and by drawing inspiration from previously parameterized compounds available in public databases (38, 62). This traditional approach presents several drawbacks: (1) it requires users to be knowledgeable about the Martini 3 force field and its intricacies; (2) it may require users to test multiple mapping schemes without any a priori guarantee of which one is optimal; (3) it depends on users knowing how to define appropriate bonded interaction networks, including the use of virtual sites, without causing simulation instability.

In this paper, we re-introduce a well-known automated algorithm (60) that provides initial models for small molecules and fragments by selecting an optimal mapping scheme and bead assignment using a built-in energy function. Previously only available for Martini 2 models, the code has now been updated to Martini 3 and includes additional improvements over the earlier implementation. We demonstrate that this new Auto-Martini version produces Martini 3 models that are stable in simulations for the large majority of tested small molecules and that reproduce transfer free energies between organic solvents and water in line with experimental logP values. Additionally, the algorithm is robust, efficient (average calculation time of 30 seconds), and user-friendly.

A key concern with automated parameterization tools for Martini is the accuracy of the generated models. Accuracy in this context refers to how well the models reproduce: (1) the AA molecular shape, (2) the bonded parameter distributions from AA simulations, and (3) thermodynamic observables like transfer free energies and even binding affinities, which are typically used for benchmarking. We first assess model quality by testing their numerical stability during CG-MD simulations and then comparing their molecular volume and shape (via SASA) to atomistic models and manually created CG models. As shown in **Figure 3**, the current implementation faithfully reproduces the molecular volume, slightly underestimating the SASA values. It also reproduces water-octanol, water-chloroform, and water-hexadecane transfer free energies with good accuracy (MAE = 2.33, 3.09, and 3.57 kJ/mol, respectively). While expert-generated models are more accurate (38), Auto-Martini achieves good accuracy at a fraction of the time needed for manual parametrization. Compared to the previous Auto-MartiniM2 (Martini 2-based), the new implementation performs better in both SASA and transfer free-energy predictions. This improvement is partly due to the upgraded Martini 3 force field, which introduces additional bead types and smaller bead sizes. This is especially relevant for capturing ring-shaped molecules like benzene, where tiny beads and accurate mapping rules are critical.

A second layer of validation involves comparing potential of mean force (PMF) profiles across membranes. Seven molecules (including four from the original Auto-Martini paper) are re-parameterized and tested in Martini 3. The resulting PMFs are compared to AA and MartiniM2 PMFs. From **Figure 4**, it appears that Martini 2 and Martini 3 models both perform reasonably well, predicting correctly the position of the main global minimum, with average errors around 1.5 kcal/mol and 1.3 kcal/mol at the water/membrane interface and interleaflet regions, respectively—an acceptable deviation. These results support the conclusion that the Auto-MartiniM3 algorithm produces models of acceptable quality for capturing key physicochemical properties, considering AA models can also vary substantially on these predictions.

To showcase the applicability of Auto-MartiniM3 for studying protein-ligand interactions, we study the binding of caffeine to the A2A receptor (**Figure 5**). This protein is a critical target in neurological drug discovery, and caffeine is a known modulator. Our CG-MD simulations showed reversible binding and unbinding events of caffeine, covering a range of residence times. Multiple entry pathways are detected, all consistent with known experimental and simulation results. The ligand found its canonical pocket, further confirming that the Auto-MartiniM3-generated caffeine model behaves as expected. This supports the potential utility of Auto-MartiniM3 in early-stage drug screening pipelines or in pocket detection in membrane environments (50).

Auto-MartiniM3 thus shows strong potential for high-throughput applications, from fragments to small molecules (up to ∼375 Da, corresponding to approximately 25 non-hydrogen atoms). It is a powerful starting point, suitable for further refinement and optimization. While the algorithm handles most small molecules robustly, specific edge cases — such as large or highly symmetric functional groups — may still result in suboptimal bead assignments or occasional instability during dynamics. In such cases, minor manual tweaks or bead adjustments can significantly enhance model performance.

Some limitations remain, including the inability to handle larger compounds efficiently — mainly due to the complexity of the underlying Voronoi-based mapping — and the restricted chemical space covered by ALOGPS-based logP predictions, particularly for underrepresented functional groups such as nitro or phosphate moieties. To extend coverage, we implement a customizable SMILES-to-logP file to improve bead-type assignment in regions where ALOGPS is known to underperform. It is worth to note, that SMILES strings are not inherently canonical. A single molecular structure can be encoded by multiple valid SMILES, potentially introducing ambiguity for structure-based analysis (91) and the quality of mapping. To circumvent this, we introduce a flag which transforms SMILES into a canon format implemented in RdKit. In addition, to improve the bonded parameters—especially for more complex molecules or when simulation instabilities arise—Auto-MartiniM3 offers a flag to generate input files compatible with Bartender (58), allowing users to refine bonded parameters in a semi-automated fashion. While user-defined mappings are not yet supported, incorporating this flexibility remains a possible future extension. By addressing these current limitations—including functional group recognition, system size scalability, and improved bonded parameter refinement— Auto-MartiniM3 can evolve into an even more robust and comprehensive tool, suitable for a broader range of applications in molecular simulations for the modeling community.

## Acknowledgments

M.S. acknowledges financial support from CALIXAR Eurofins for her doctoral research project. T.B. and L.J.W. acknowledge support by the Deutsche Forschungsgemeinschaft (DFG, German Research Foundation) under Germany’s Excellence Strategy EXC 2181/1 - 390900948 (the Heidelberg STRUCTURES Excellence Cluster) and the Klaus Tschira Foundation (SIMPLAIX). G.P.P. and P.C.T.S. acknowledge the support provided by the CNRS and PharmCADD. They also acknowledge the support of the Centre Blaise Pascal’s IT test platform at ENS de Lyon (Lyon, France) for the computer facilities. The platform operates the SIDUS solution developed by Emmanuel Quemener (Quemener and Corvellec, 2013). This work was granted access to the HPC resources of CALMIP supercomputing center (under the allocation P24033) and TGCC Joliot-Curie supercomputer (under the GENCI allocations A0180716209, A0160715135, and SS010715367).

