## Supplementary Information for "Fast parameterization of Martini3 models for fragments and small molecules"

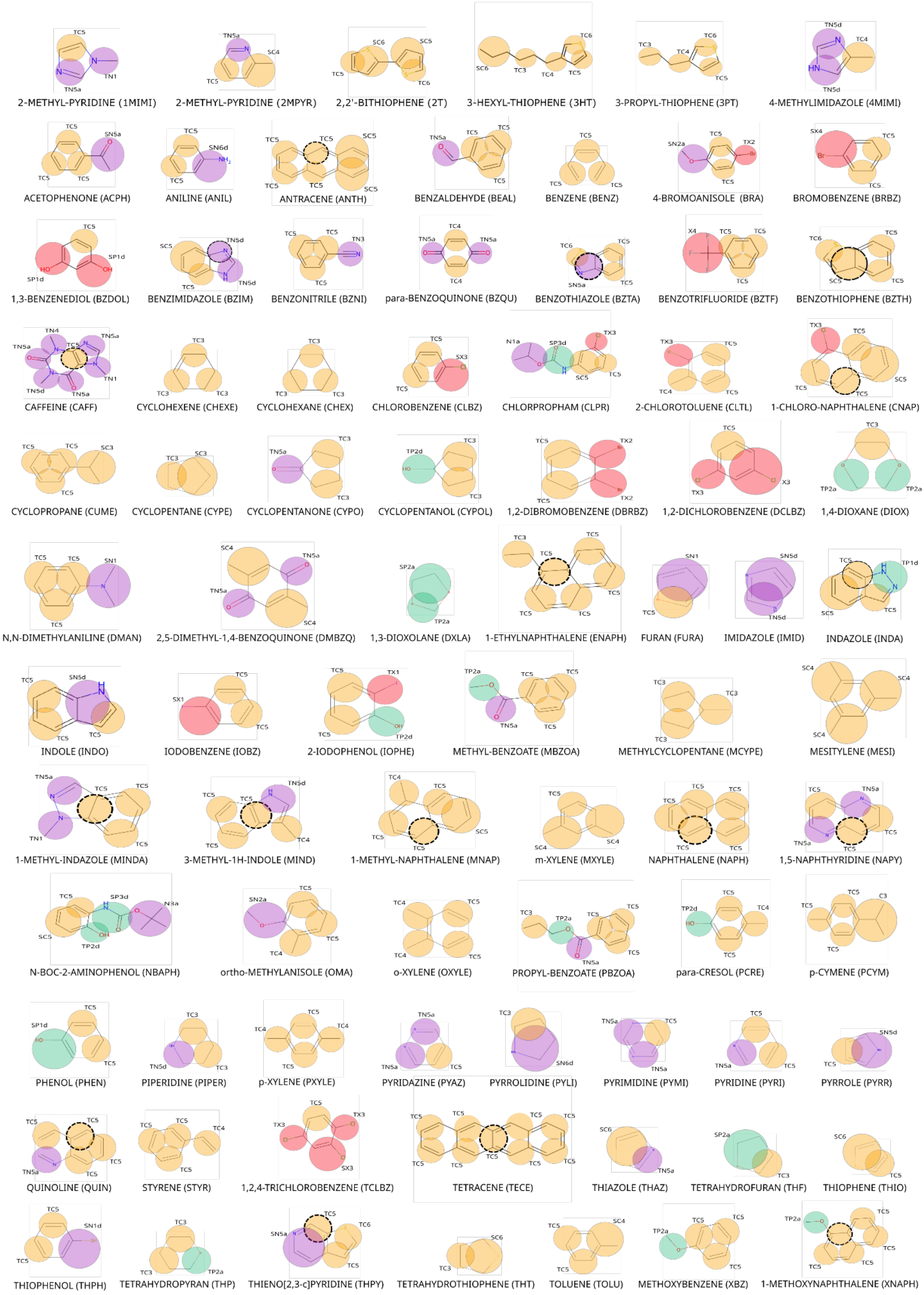


**Figure S1:** CG-mapping of molecules extracted from Martini Database (62) made with Auto-MartiniM3.

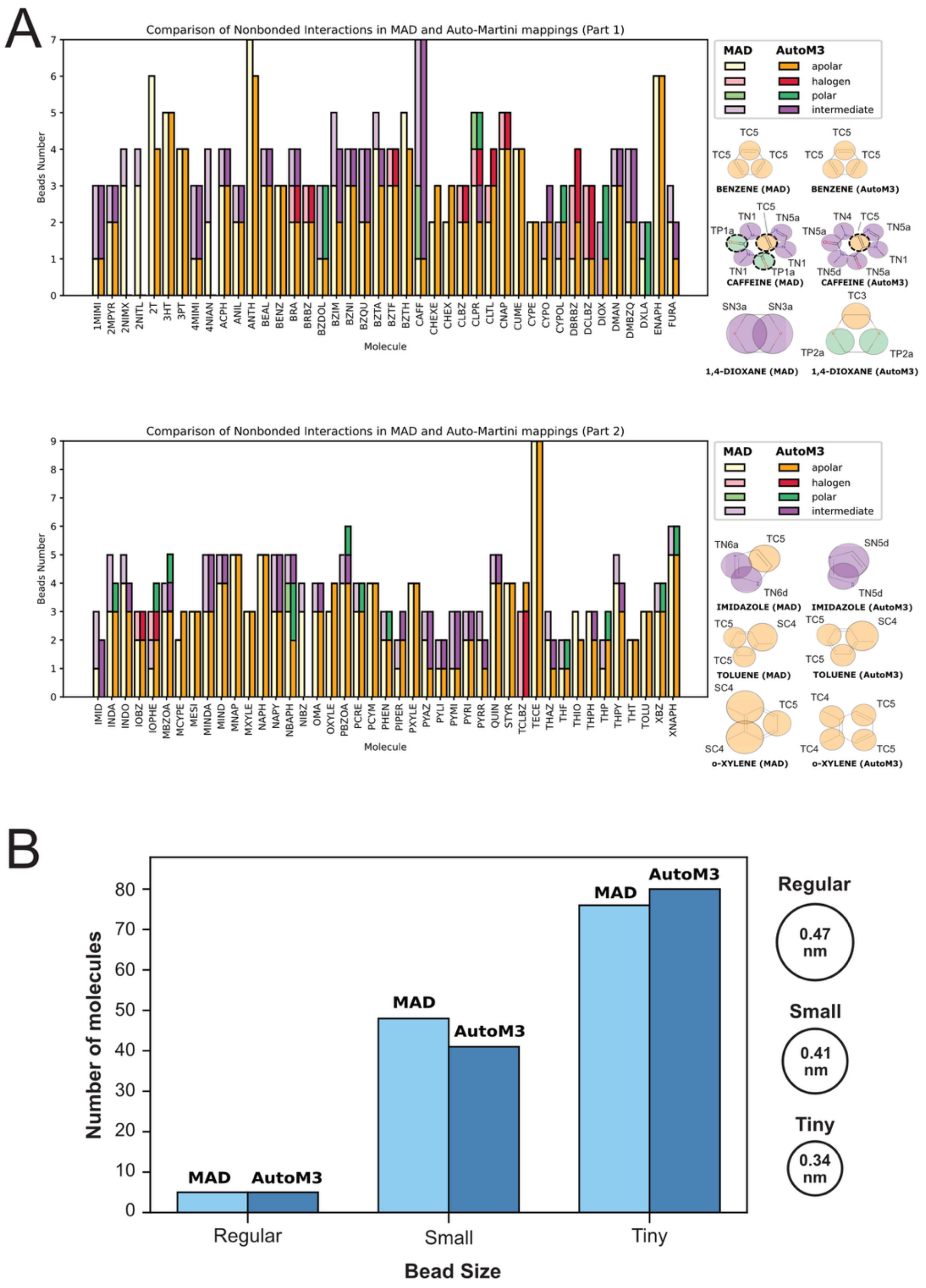


**Figure S2: Comparison of topology files created by Auto-MartiniM3 and Martini3 developers. (A)** Models parametrizations made by Martini 3 force field developers, available in Martini Database (MAD) (62) and parametrizations created automatically by Auto-MartiniM3. The number of beads from all models are represented by barplot height and bead types are differentiated by colors, apolar beads in orange, ion beads in blue, polar beads in green and intermediate polarity beads in pink. Light colors (left bars) represent data from MAD and dark colors (right bars) represent data from AutoM3 models. Missing bars corresponds to models that Auto-martiniM3 did not manage to parametrize. **(B)** Number of models containing regular, Small, and Tiny beads for MAD (light blue) and for Auto-MartiniM3 (dark blue).

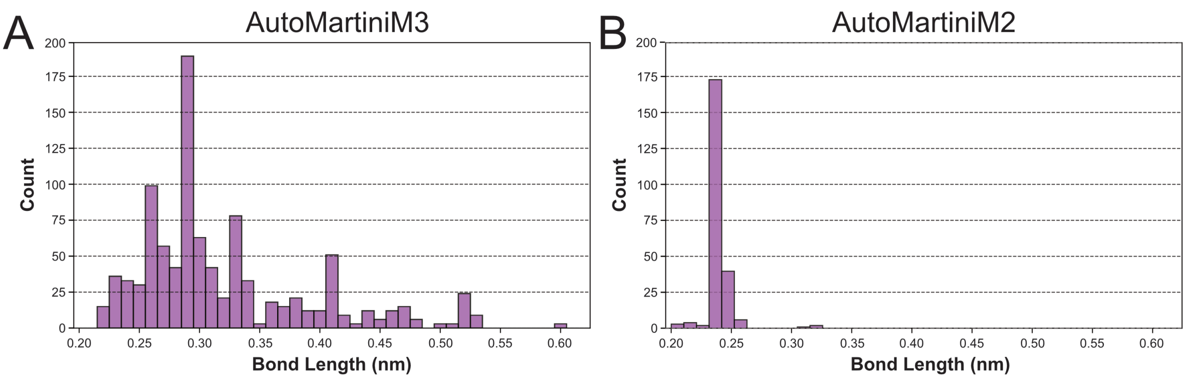


**Figure S3:** Distribution of bond and constraints length parametrized by Auto-MartiniM3 **(A)** and Auto-MartiniM2 **(B)**.

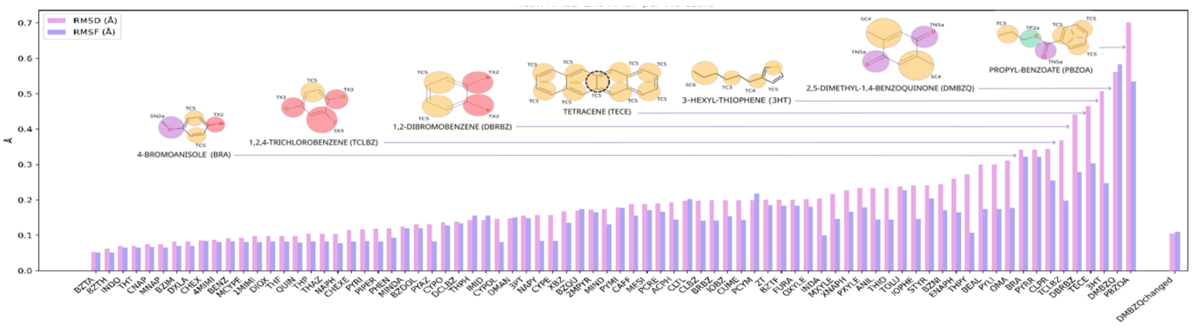


**Figure S4:** Mean RMSD / RMSF of Auto-MartiniM3 models: RMSD (purple) and RMSF (blue) data with few, more dynamic molecules depicted above. DMBZQ model with increased force values for bonded parameters is depicted on the right of the plot as “DMBZQchanged”.

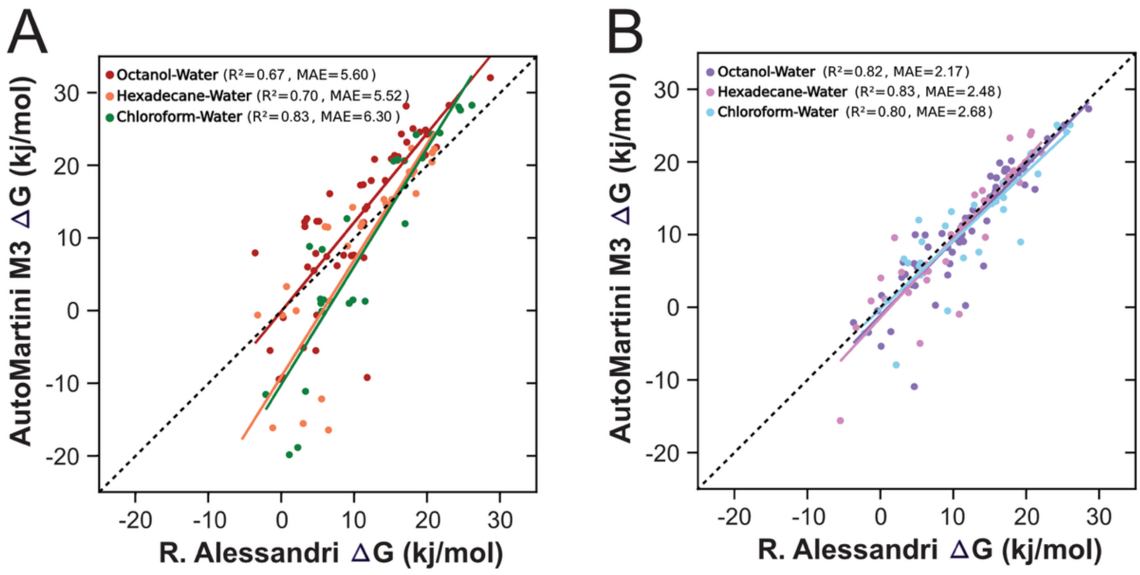


**Figure S5:** Partitioning Behaviour of original Auto-MartiniM2 **(A)** and Auto-MartiniM3 **(B)** in water/oil phases in regard to R.Alessandri data (38) .

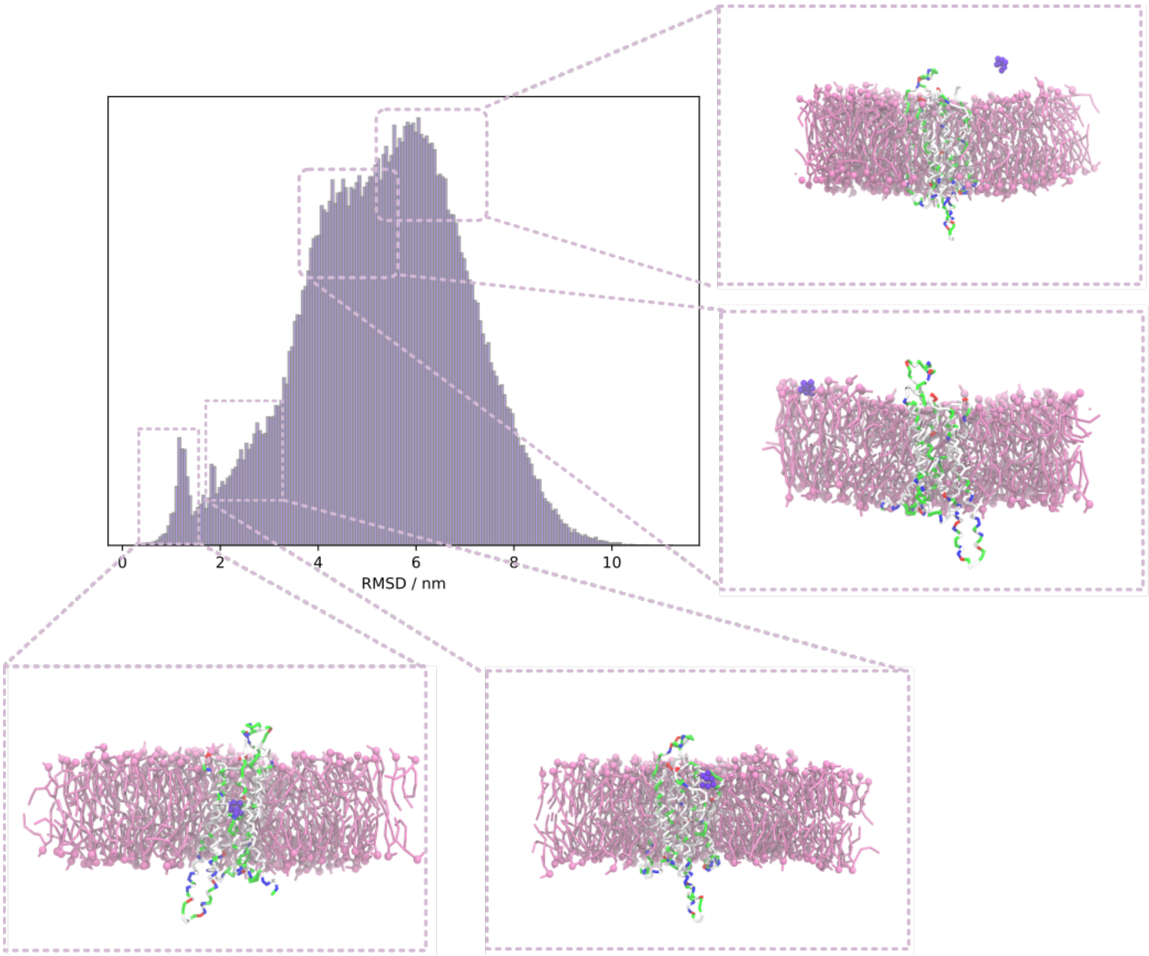


**Figure S6:** RMSD of Auto-MartiniM3-generated caffeine's example positions in binding site to A2A receptor during a 20 microsecond simulation, with coarse-grained crystalline structure of the system as a reference.

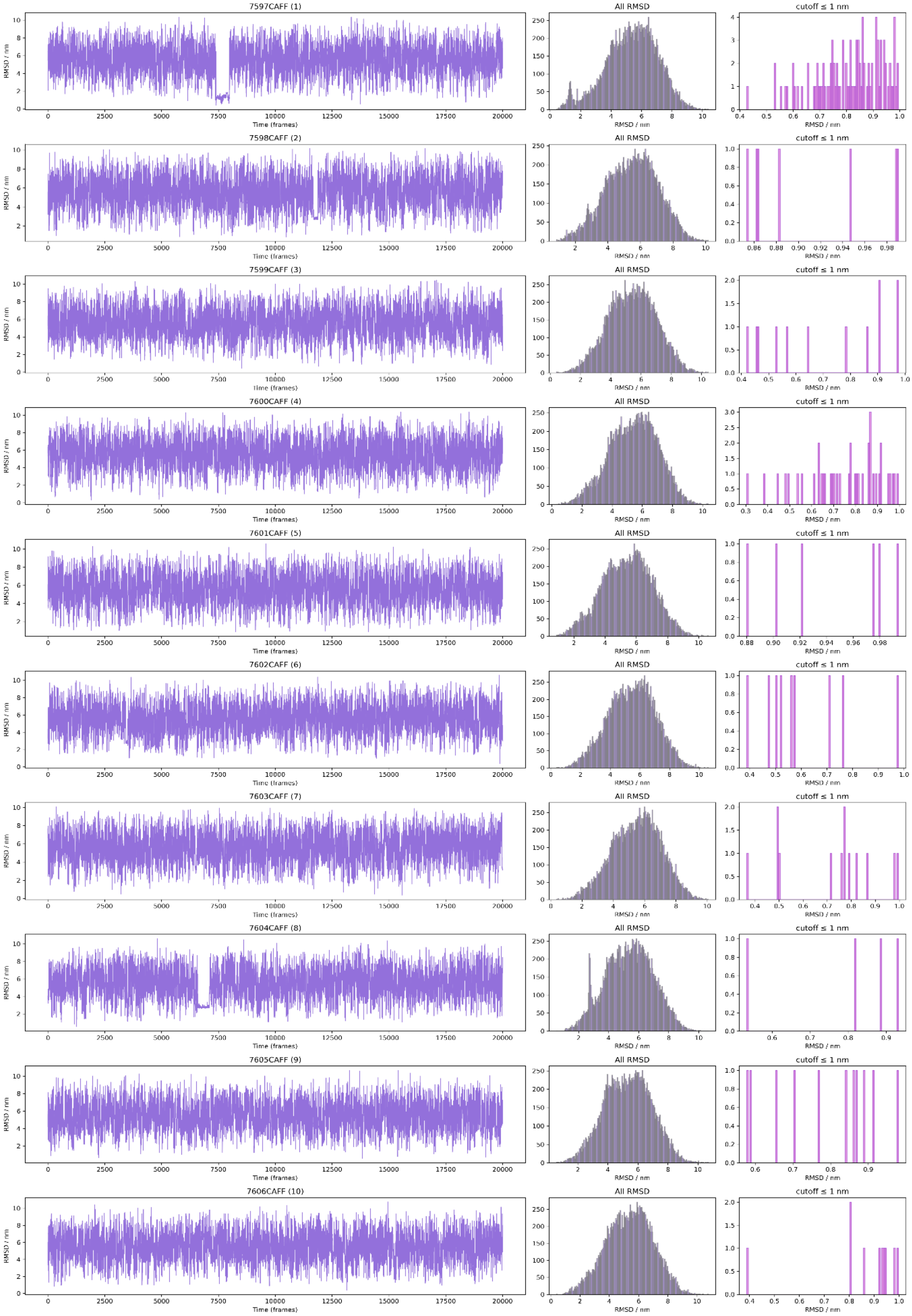


**Figure S7.1:** RMSD of Auto-MartiniM3-generated caffeine in reference to caffeine in its binding site seen in Xray data (Simulation 1).

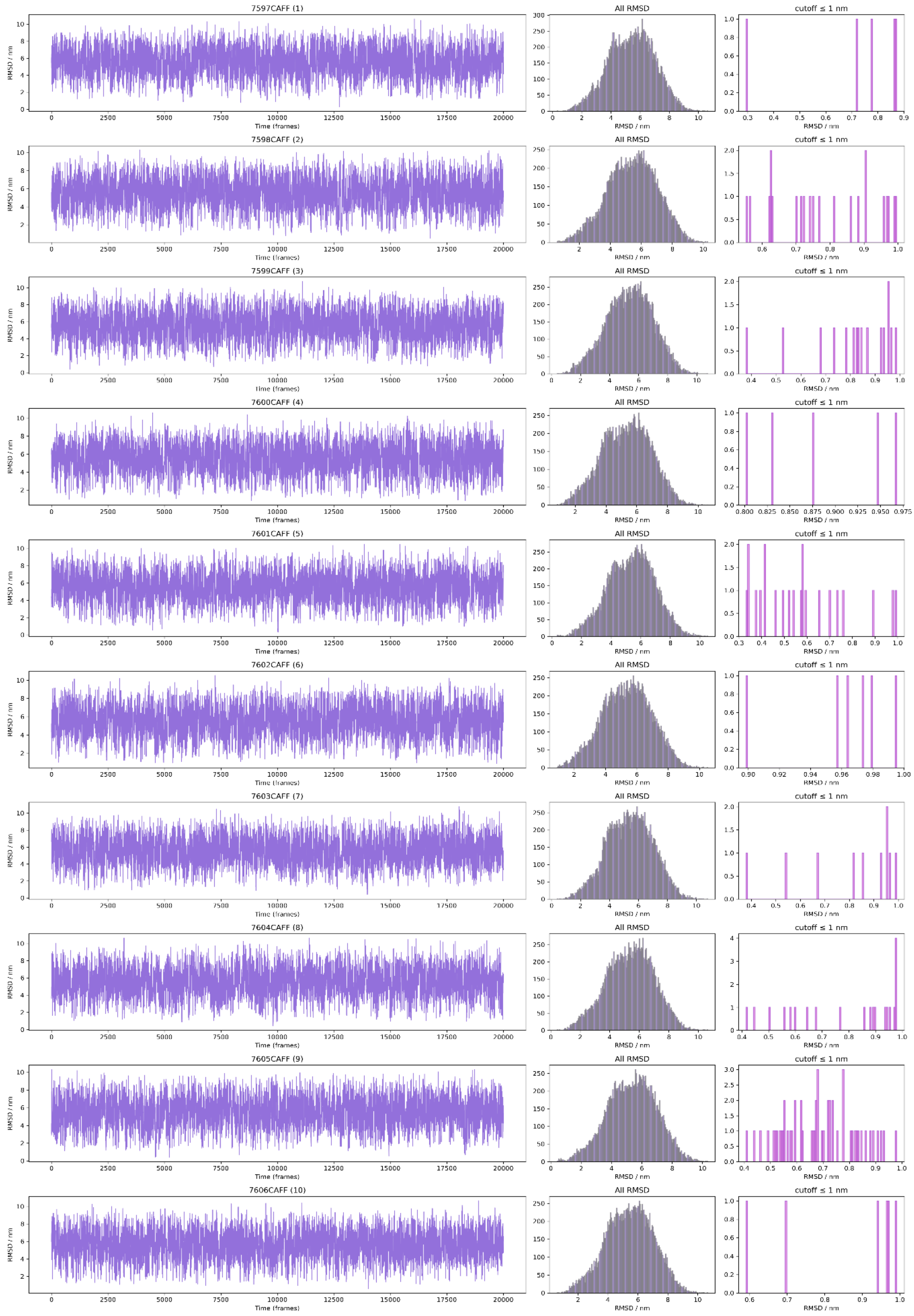


**Figure S7.2:** RMSD of Auto-MartiniM3-generated caffeine in reference to caffeine in its binding site seen in Xray data (Simulation 2).

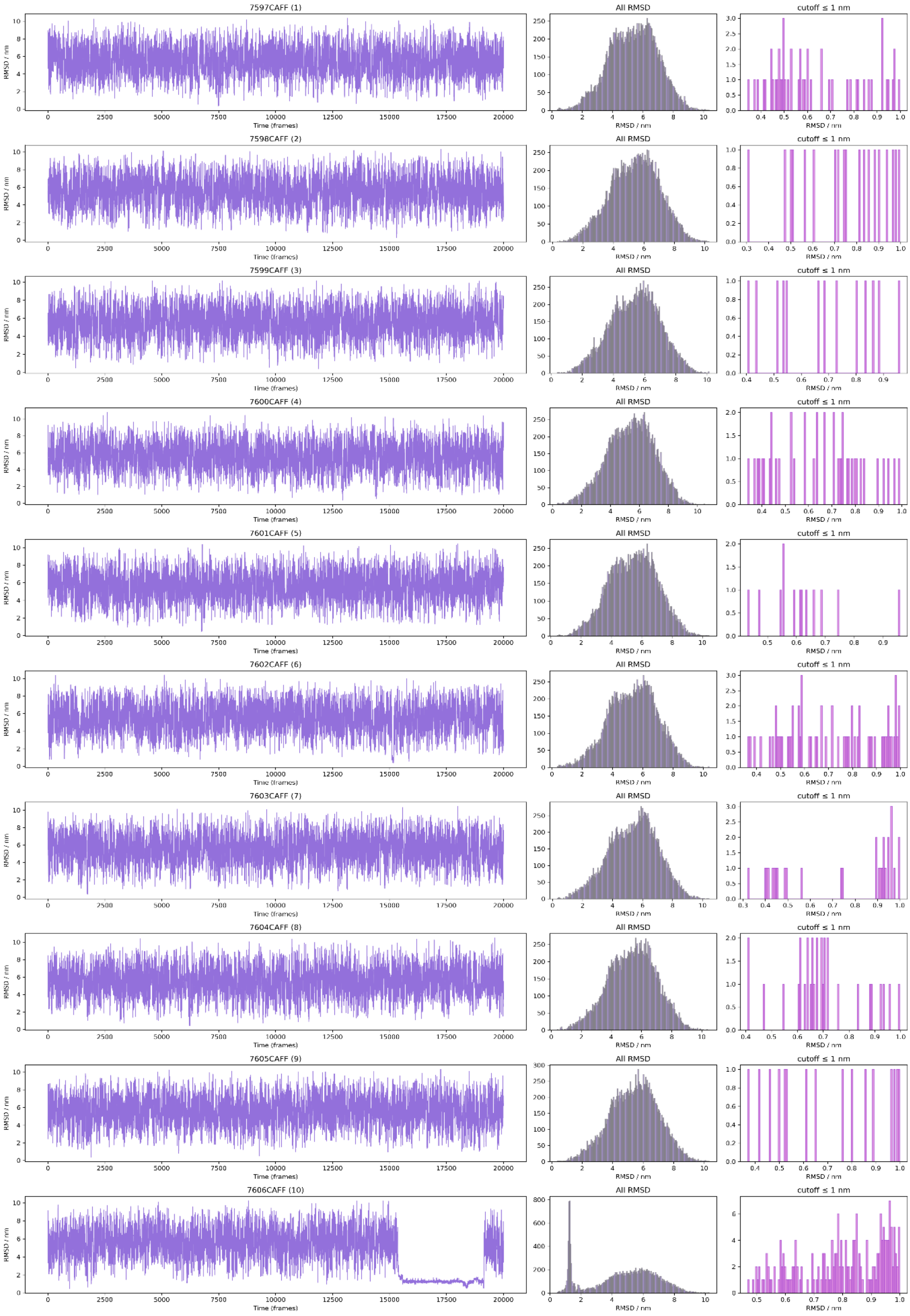


**Figure S7.3:** RMSD of Auto-MartiniM3-generated caffeine in reference to caffeine in its binding site seen in Xray data (Simulation 3).

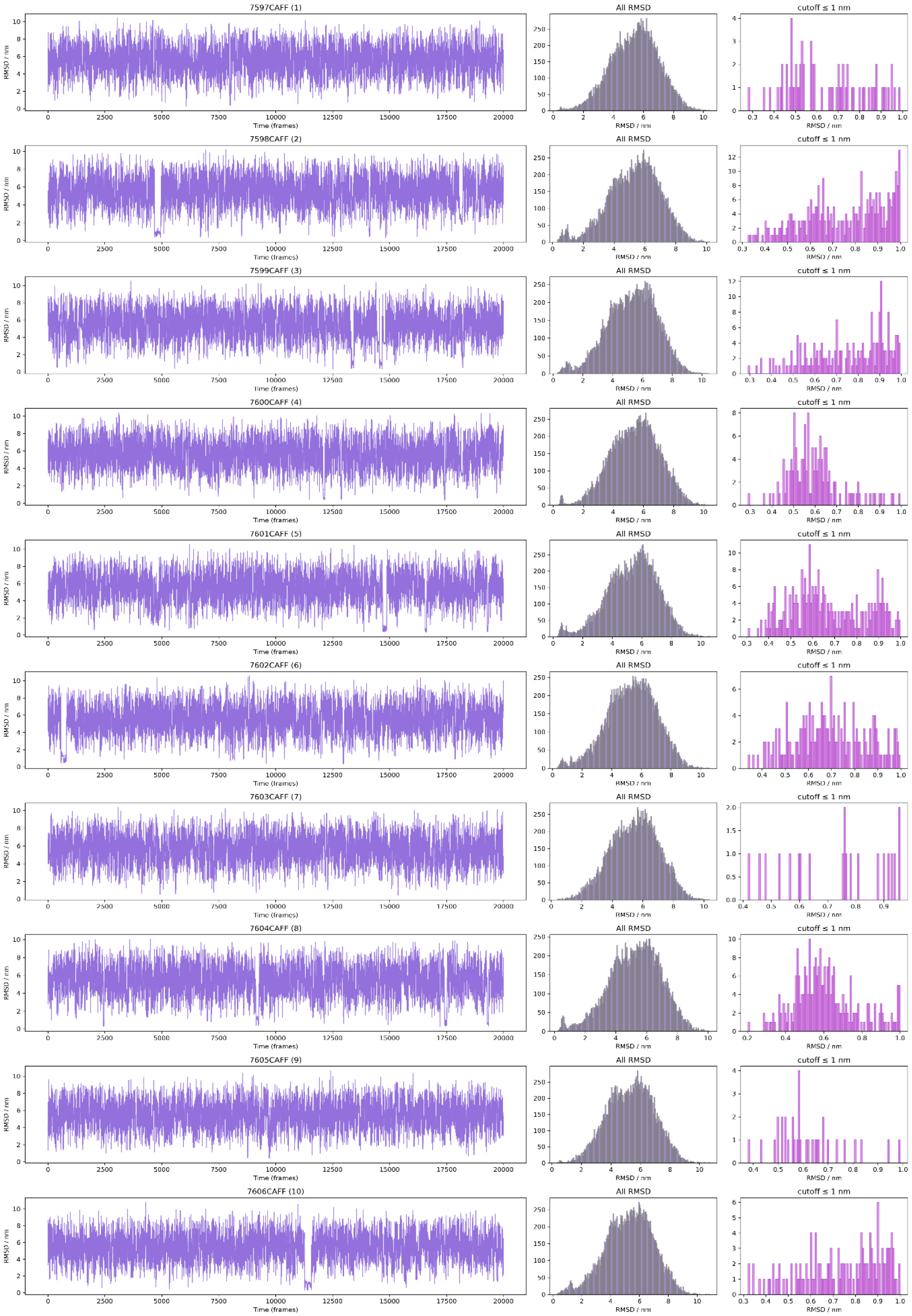


**Figure S7.4:** RMSD of Auto-MartiniM3-generated caffeine in reference to caffeine in its binding site seen in Xray data (Simulation 4).

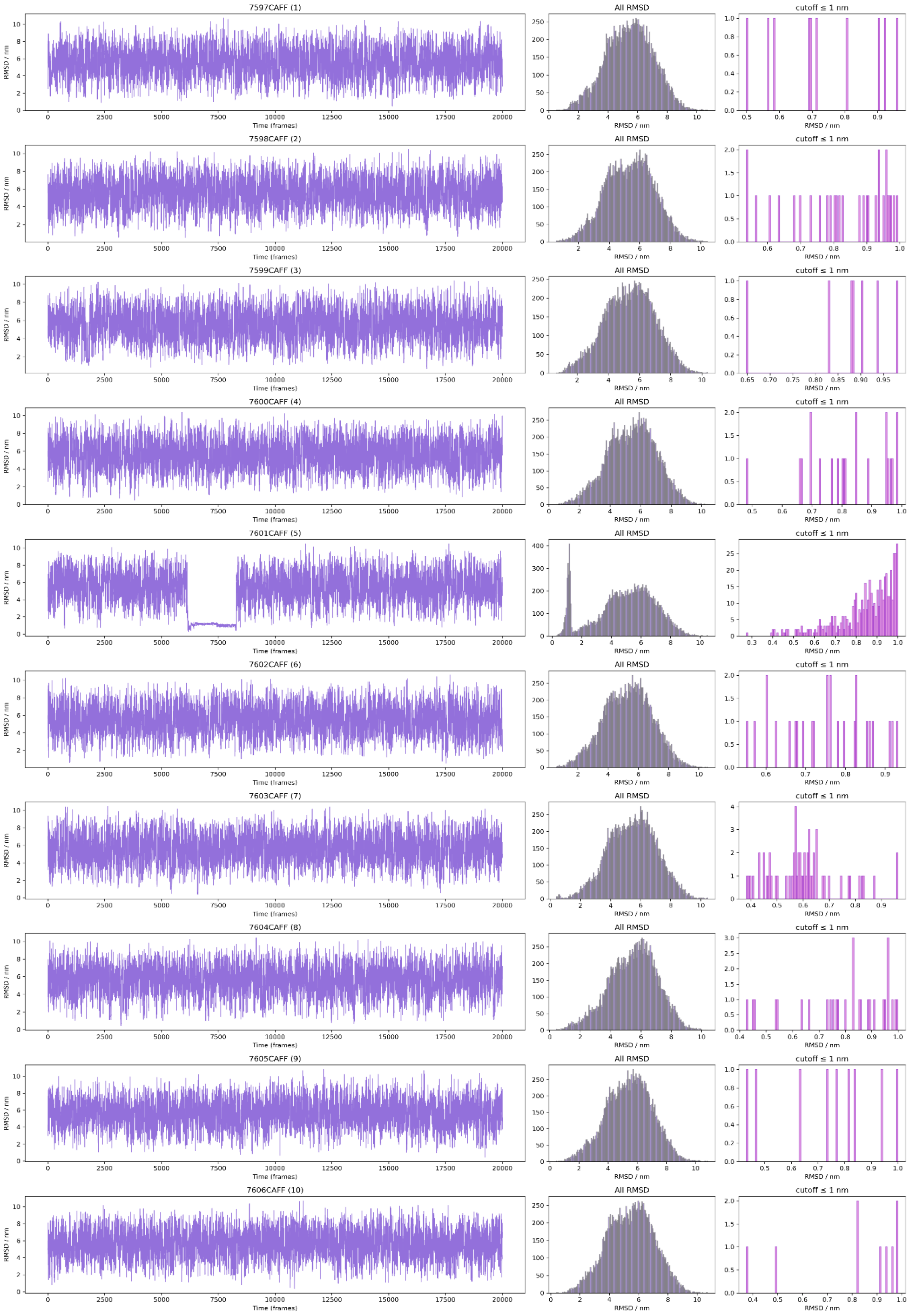


**Figure S7.5:** RMSD of Auto-MartiniM3-generated caffeine in reference to caffeine in its binding site seen in Xray data (Simulation 5).

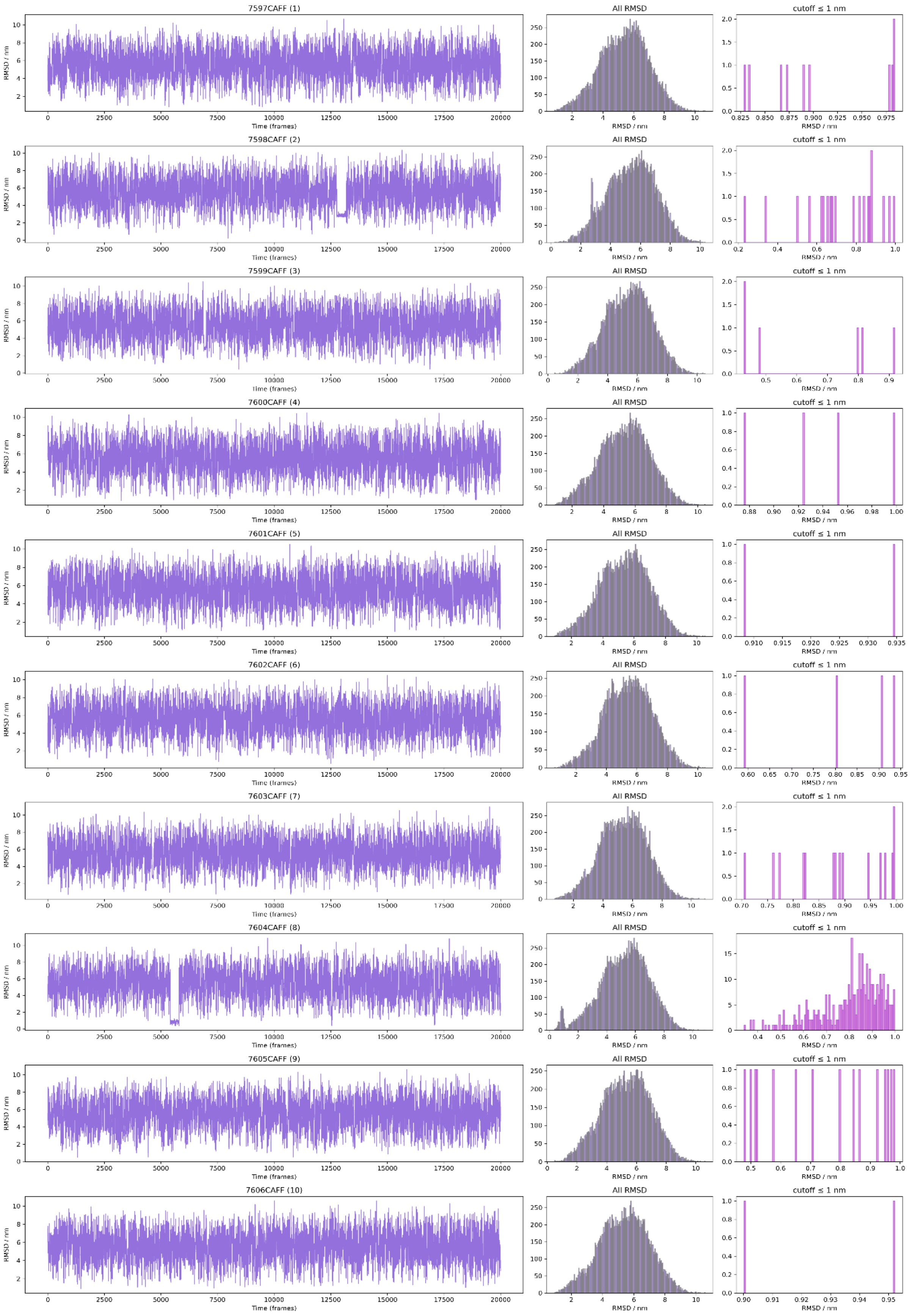


**Figure S7.6:** RMSD of Auto-MartiniM3-generated caffeine in reference to caffeine in its binding site seen in Xray data (Simulation 6).

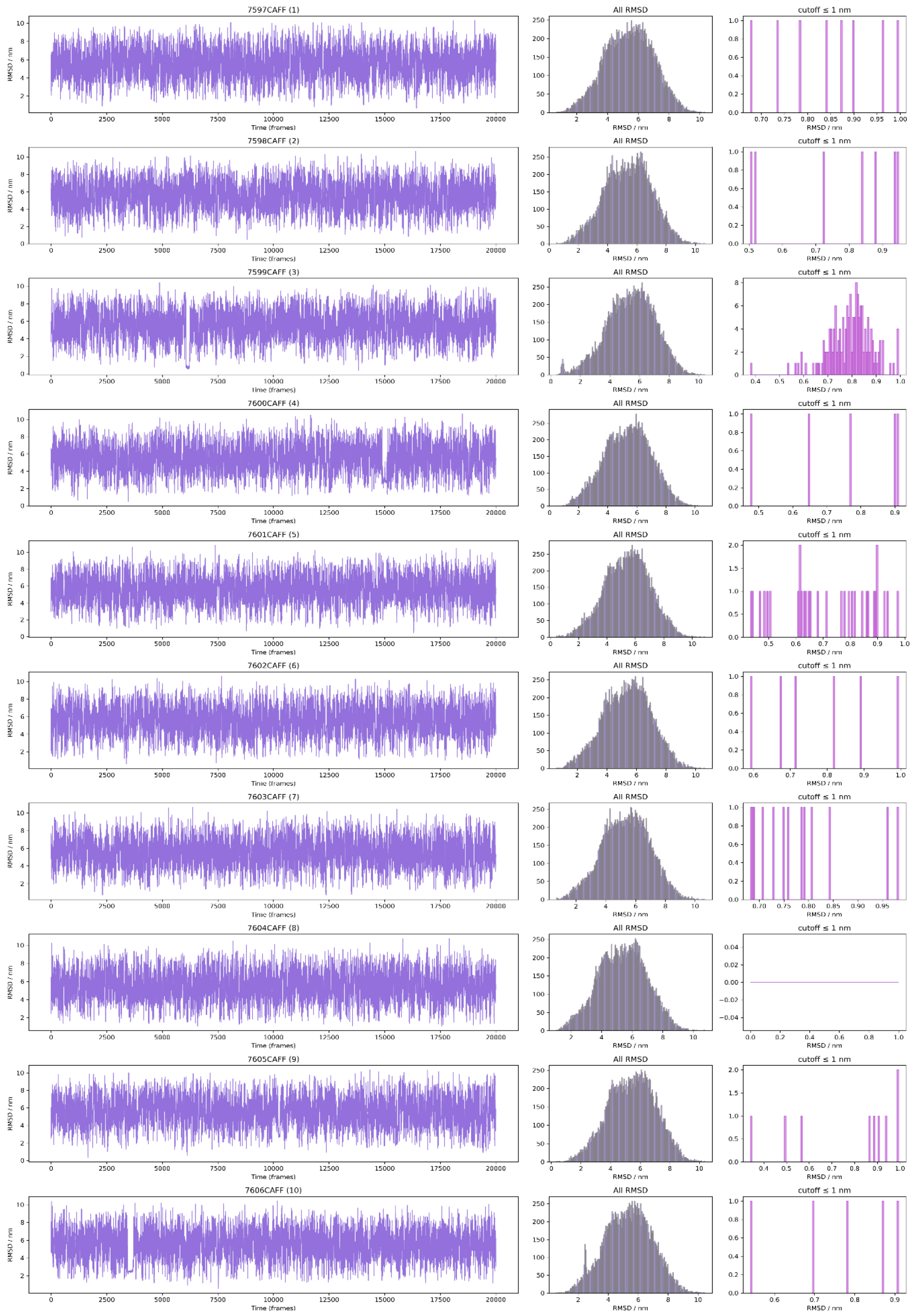


**Figure S7.7:** RMSD of Auto-MartiniM3-generated caffeine in reference to caffeine in its binding site seen in Xray data (Simulation 7).

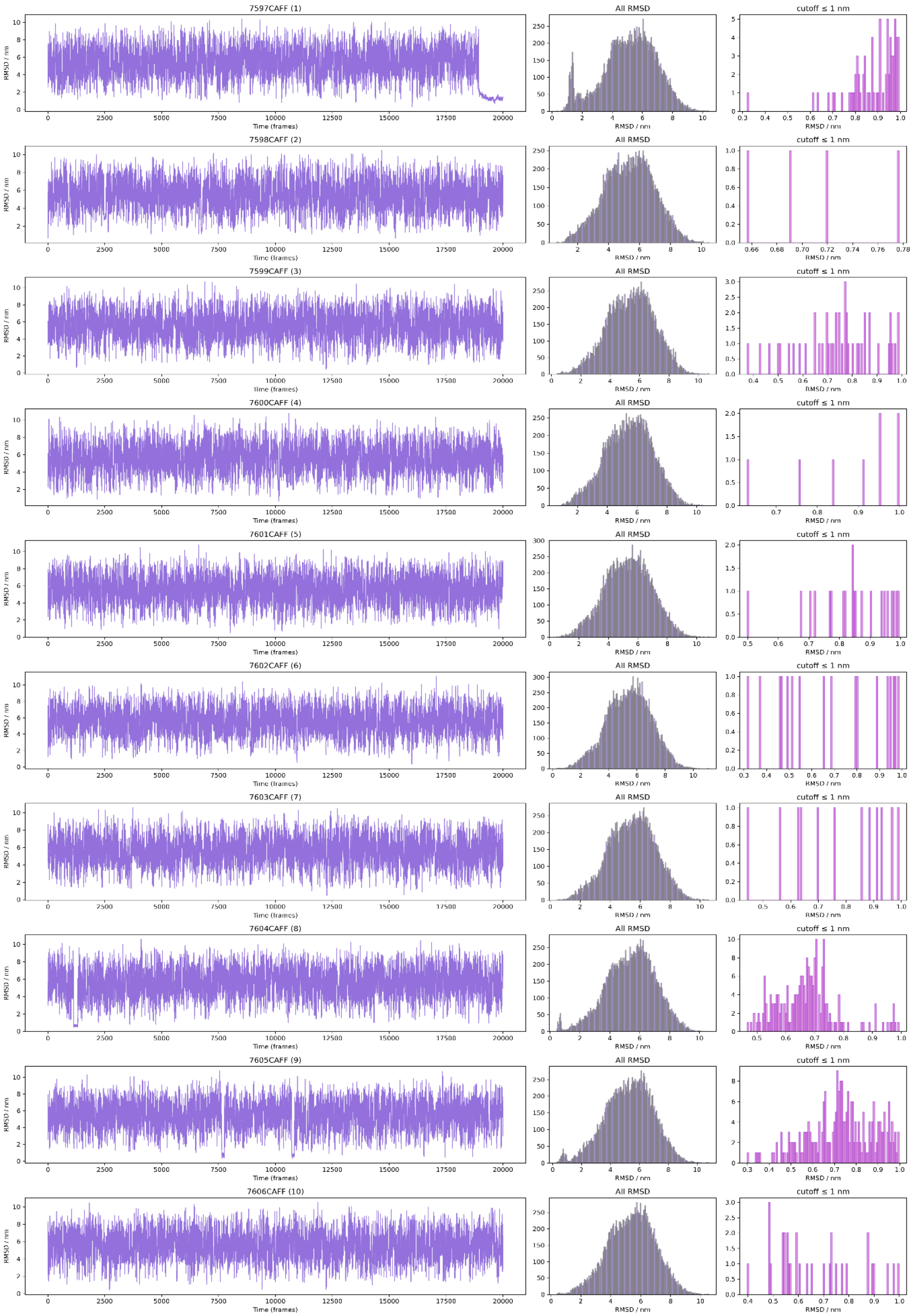


**Figure S7.8:** RMSD of Auto-MartiniM3-generated caffeine in reference to caffeine in its binding site seen in Xray data (Simulation 8).

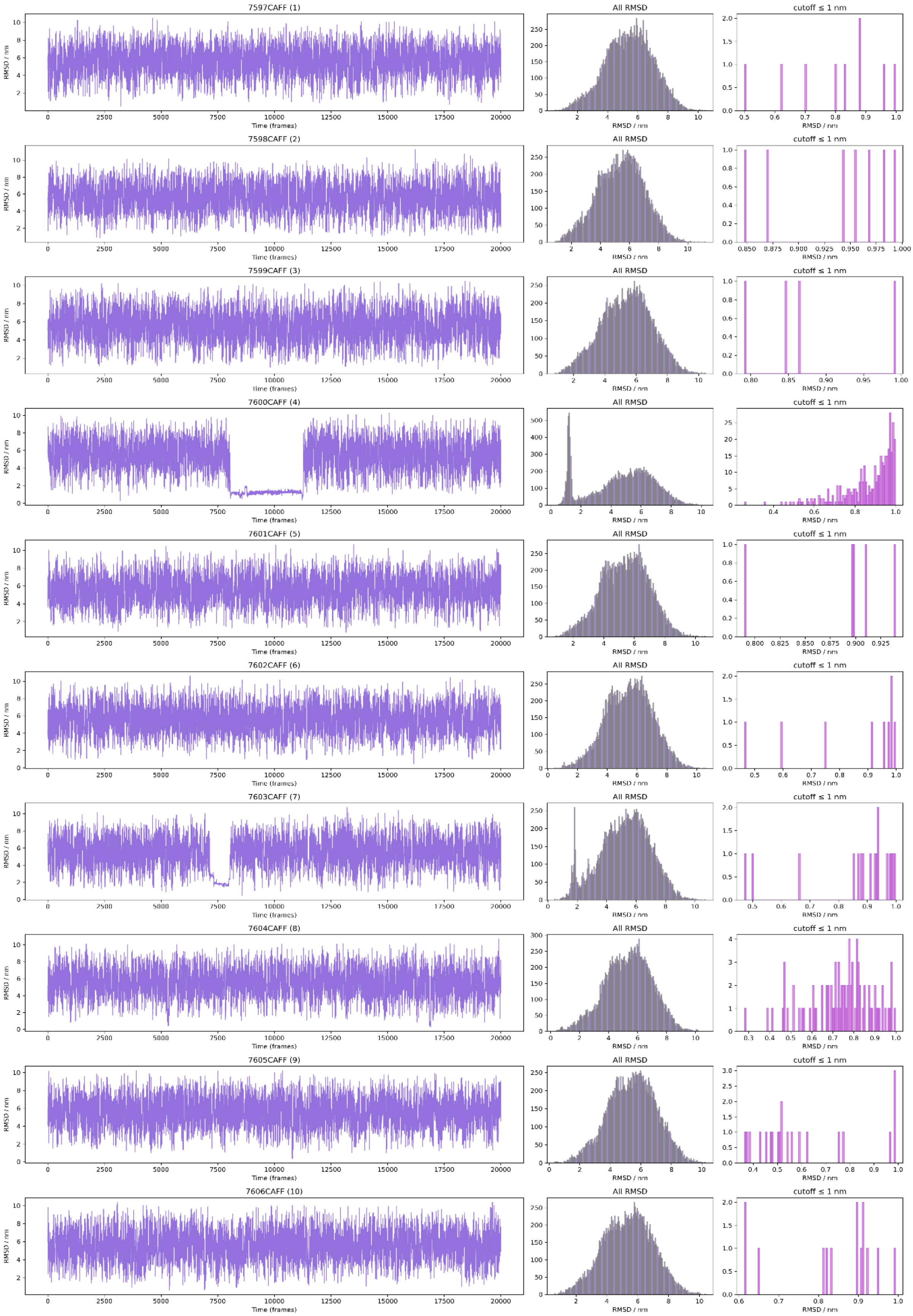


**Figure S7.9:** RMSD of Auto-MartiniM3-generated caffeine in reference to caffeine in its binding site seen in Xray data (Simulation 9).

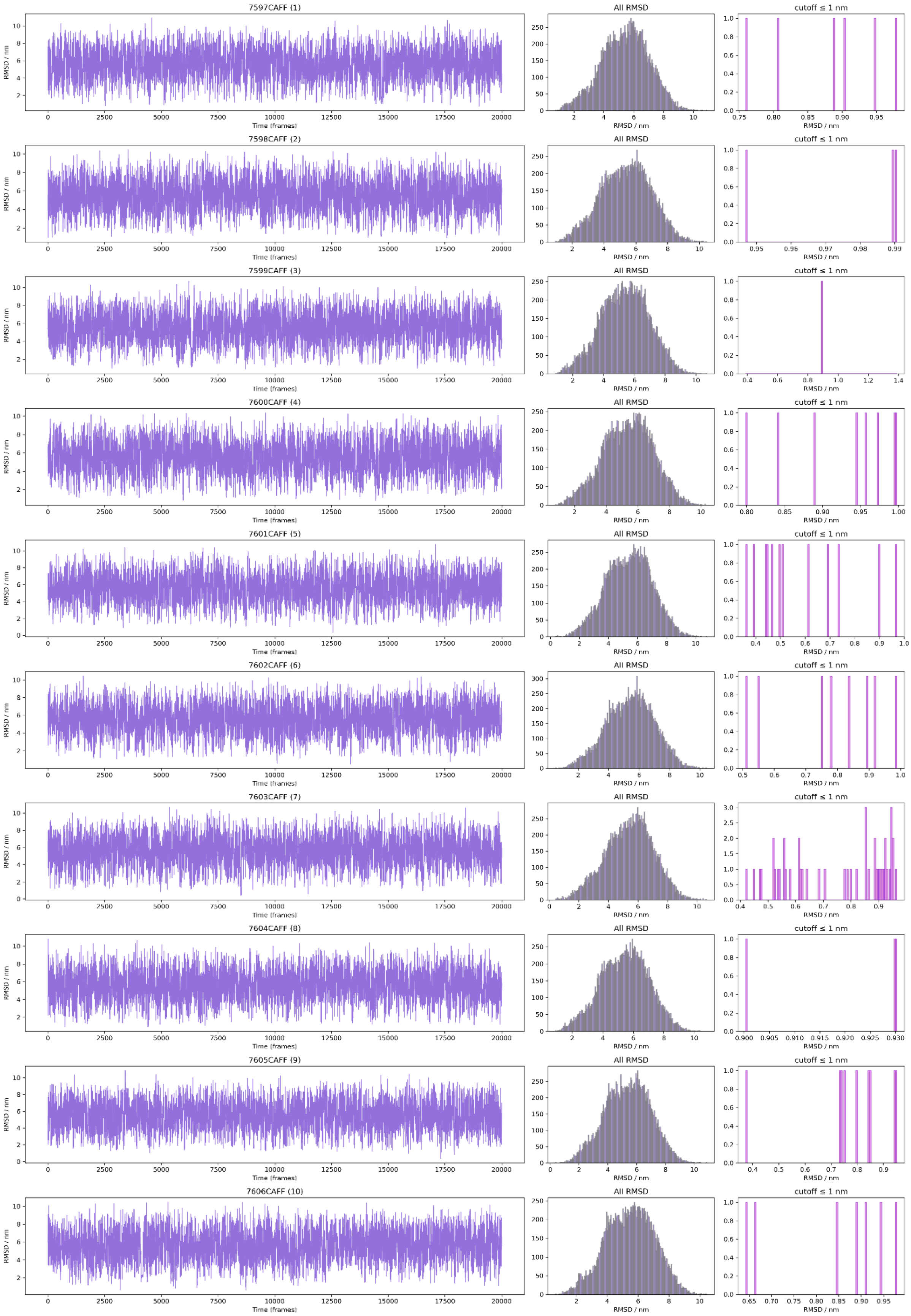


**Figure S7.10:** RMSD of Auto-MartiniM3-generated caffeine in reference to caffeine in its binding site seen in Xray data (Simulation 10).

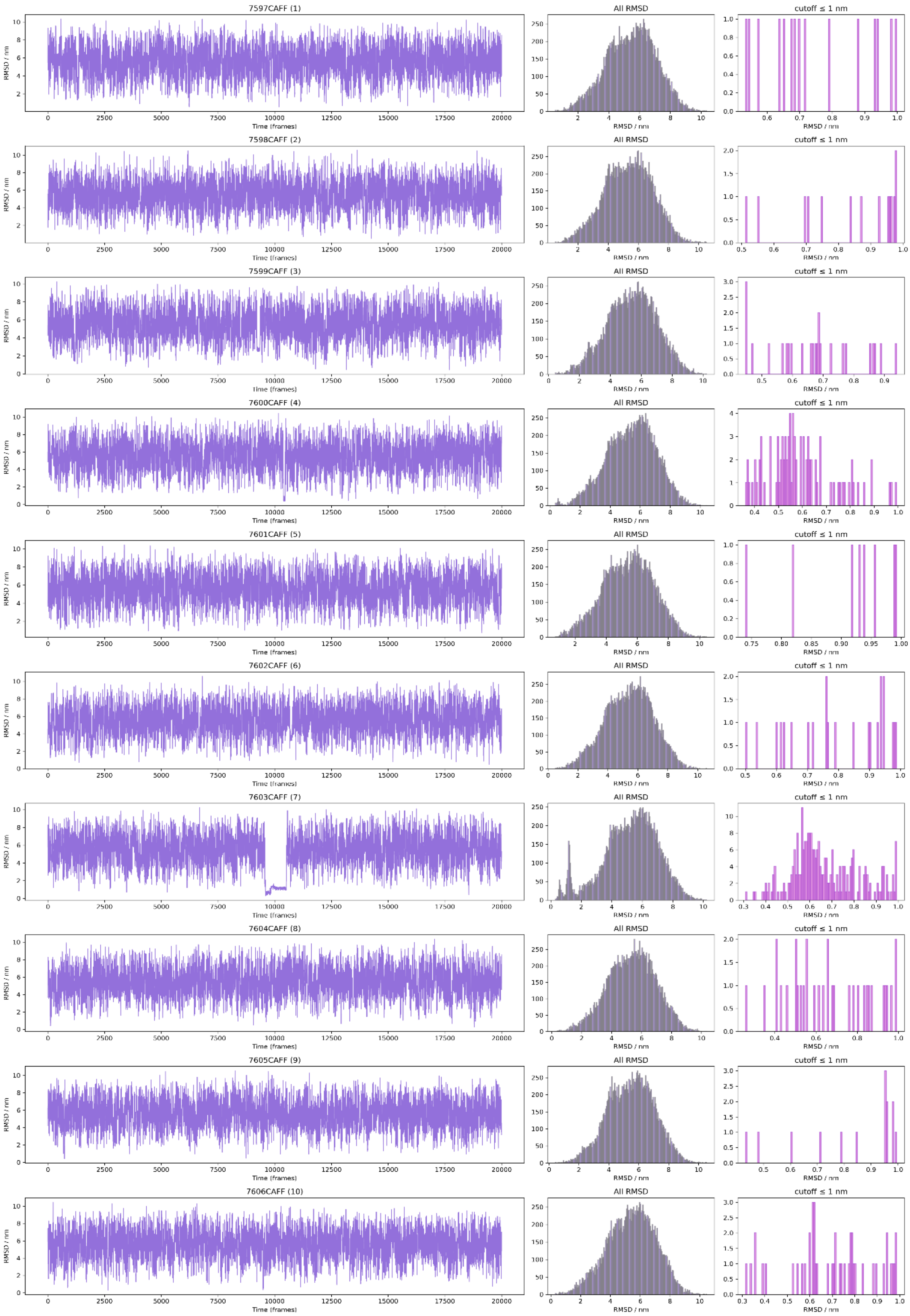


**Figure S7.11:** RMSD of Auto-MartiniM3-generated caffeine in reference to caffeine in its binding site seen in Xray data (Simulation 11).

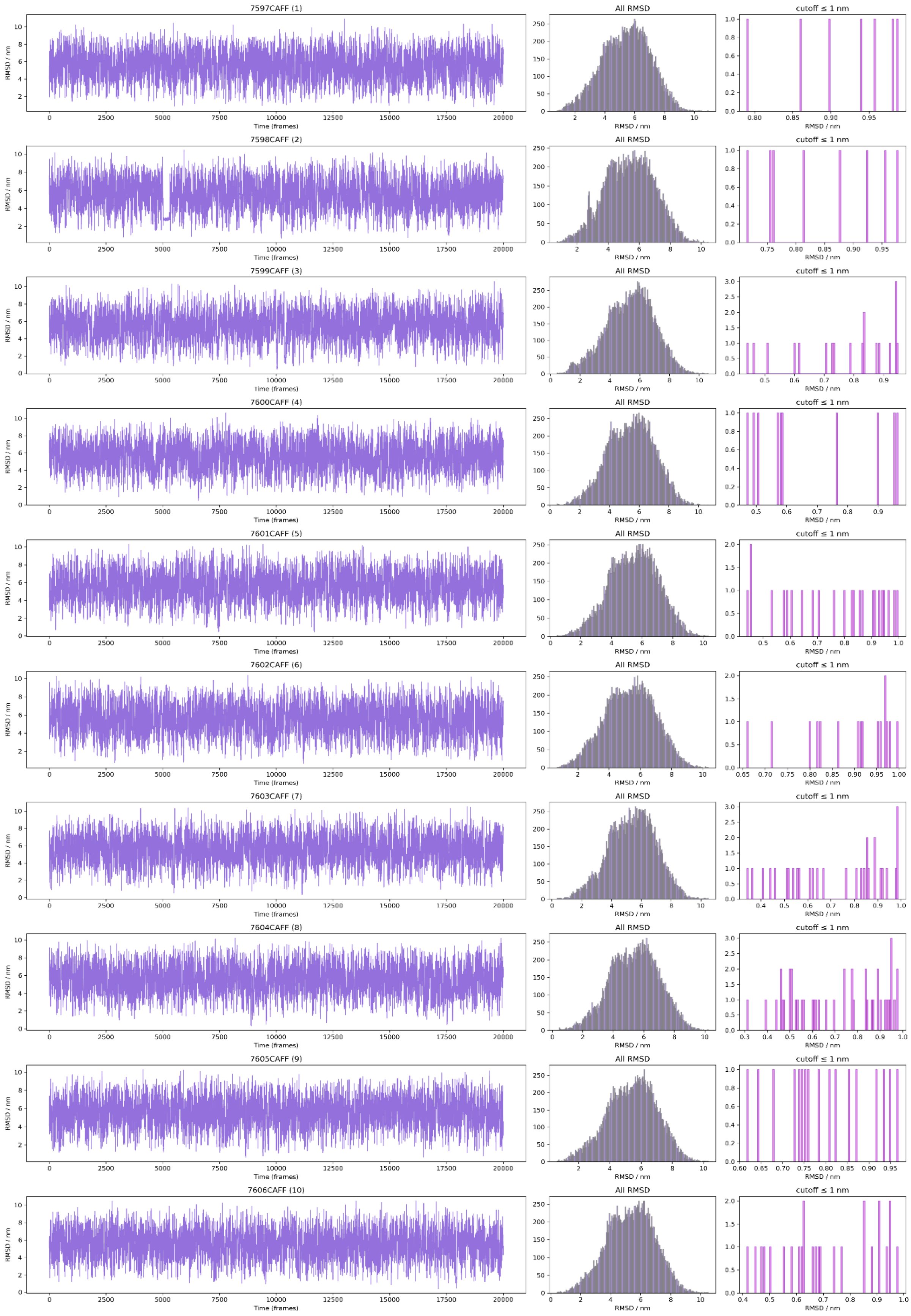


**Figure S7.12:** RMSD of Auto-MartiniM3-generated caffeine in reference to caffeine in its binding site seen in Xray data (Simulation 12).
